# Normalization by Valence and Motivational Intensity in the Sensorimotor Cortices (PMd, rM1, and cS1)

**DOI:** 10.1101/702050

**Authors:** Zhao Yao, John P Hessburg, Joseph Thachil Francis

## Abstract

Our brain’s ability to represent vast amounts of information, such as continuous ranges of reward spanning orders of magnitude, with limited dynamic range neurons, may be possible due to normalization. Recently our group and others have shown that the sensorimotor cortices are sensitive to reward value. Here we ask if psychological affect causes normalization of the sensorimotor cortices by modulating valence and motivational intensity. We had two non-human primate (NHP) subjects (one male bonnet macaque and one female rhesus macaque) make visually cued grip-force movements while simultaneously cueing the level of possible reward if successful, or timeout punishment, if unsuccessful. We recorded simultaneously from 96 electrodes in each the following: caudal somatosensory, rostral motor, and dorsal premotor cortices (cS1, rM1, PMd). We utilized several normalization models for valence and motivational intensity in all three regions. We found three types of divisive normalized relationships between neural activity and the representation of valence and motivation, linear, sigmodal, and hyperbolic. The hyperbolic relationships resemble receptive fields in psychological affect space, where a unit is susceptible to a small range of the valence/motivational space. We found that these cortical regions have both strong valence and motivational intensity representations.

## Introduction

Neural networks within the primary sensorimotor cortices need to function within a limited dynamic range while encoding multiple forms of information such as kinematics and kinetics (Georgopoulos et al., 1982, 1992; Li et al., 2001; Chhatbar and Francis, 2013). The nervous system could, in part, increase its information-carrying capacity by utilizing a temporal code, which would be less sensitive to a limited dynamic range than a rate code (Bialek et al., 1996). Another possibility is provided by divisive normalization (Carandini and Heeger, 2011), where the input from another region and/or from the local population can dynamically normalize the network, so neural responses remain within a finite range. Such divisive normalization has been suggested as a canonical neural computation, and has been seen in neural systems from primary sensory cortical areas (Heeger, 1992; Schwartz and Simoncelli, 2001; Sato et al., 2016), regions involved in multisensory integration (Ohshiro et al., 2017) to networks involved in decision-making (Louie et al., 2011, 2014; Gluth et al., 2020; Webb et al., 2020).

Recently, it has been shown that the primary motor (M1) and somatosensory (S1) cortices encode reward in addition to kinematics and kinetics (Marsh et al., 2015; McNiel et al., 2016b, 2016a; Ramkumar et al., 2016; Ramakrishnan et al., 2017; An et al., 2019; Moore and Francis, 2020). However, there has been little mention of psychological affect more broadly in the neurophysiological literature on sensorimotor integration, control, and learning in S1 and M1. Affect here refers to information related to the internal representation of feelings or emotions (Rolls, 2014), also see the definition of affect as stated by the American Psychological Association (APA). Valence is the value associated with a task or object that ranges from positive to negative, reflecting stimuli ranging from positive valence (rewarding) to negative valence (punishing). These are associated with approach and avoidance behaviors, respectively. In contrast to valence, the interaction between reward and punishment can also be represented as motivational intensity or motivational salience. Here, reward and punishment would increase motivation, as individuals are driven to avoid punishment and likewise driven to obtain reward. A neutral stimulus would have zero motivation in this sense and zero valence.

While affect’s influence on many brain regions has been studied, this has not been the case in the primary sensorimotor cortices, although many tasks involving sensorimotor control may be impacted by these variables (Marsh et al., 2015; Ramkumar et al., 2016; Ramakrishnan et al., 2017; Tarigoppula et al., 2018; Zhao et al., 2018; An et al., 2019). Therefore, we characterized neural activity to cued possible reward and simultaneously cued possible punishment in cS1, rM1, and PMd. By cuing both reward and punishment, in this manner, we modulated valence and motivational intensity (Roesch and Olson, 2004).

The neural firing rates and time to complete a trial can be modeled well by divisive normalization models. Here we demonstrate that neural modulation associated with cued reward and punishment in an operant conditioning task, where reward or punishment was delivered according to an individual’s performance, is encoded in cS1, rM1, and PMd. We obtained similar results using either a divisive term based on a combined variable comprised of the cued reward level and a scaled version of the cued punishment level or simply a function of the brain region’s population activity. We found a strong representation of both valence and motivational intensity. The dominant of these two variables depends on the time in a trial, such as post-cue vs. post-feedback. Brain regions responded similarly, indicating a common driving region or signal led to our results.

## Methods

### Behavioral task and rational

In this work, we hypothesized that the sensorimotor cortices (cS1, rM1, and PMd) represent, or are at least modulated by, cued affective information. As these regions are involved in moving, we wanted to use a task with no movement by the non-human primates (NHPs) during the immediate post-affective cue period. These NHPs and this task were also used during brain-machine interfacing experiments (Zhao et al., 2018) after the data under consideration here were taken. The periods used in our analysis presented here were not during any actual movement produced by the NHP.

Two NHPs, one male rhesus macaque (NHP S, Macaca mulatta) and one female bonnet macaque (NHP P, Macaca radiata), were trained to perform an isometric grip force task (Fig. 1). In this task, subjects controlled certain aspects of a virtual anthropomorphic robotic arm (Barrett WAM) interacting with a virtual cylindrical object. Each trial consisted of 6 stages: cue display, autonomous robot reaching, manual isometric grip force output, autonomous robot transporting, manual isometric releasing, and feedback, which could be reward delivery if it was a successful trial or a timeout period if not. At the start of a trial, affective cues were displayed at the top of the virtual environment. The number of green squares indicated the level of fruit juice reward that the NHP would receive upon successful completion of the task, with the number present (0 - 3) corresponding to the number of 0.5 second juice delivery periods. If no green squares were displayed, then no reward was delivered upon successful completion of the trial. Red squares indicated the level of timeout punishment the NHP would receive if the trial was completed unsuccessfully. The number of red squares (0 - 3) corresponded to the number of 5 second timeout periods. A transparent red screen was displayed on the video monitor over the environment, and the animal had to wait for the subsequent trial to start. Unsuccessful trials were repeated at the same reward and punishment level until completed successfully. This repeated trial method motivated the subjects to successfully complete non-rewarding trials by not allowing them to purposefully fail and skip low-value trials, which they did before moving to this format.

**Figure 1.**
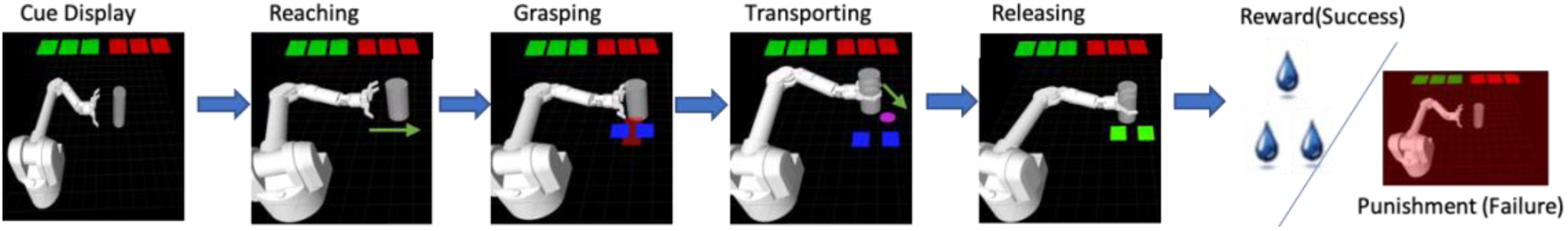
Grip Force Task Cued grip force task. The behavioral task was composed of 6 scenes for each trial as described in the text.

During our isometric grip force task, the virtual arm automatically reached a target cylinder (Fig.1). The NHP controlled the robotic grasping motion via isometric force output applied to a force transducer with its right hand. The amount of force used was represented in the virtual environment by a red rectangle that increased in width proportional to the NHPs force output and by the fingers of the robotic hand grasping the object as this force was applied. The subject had to apply, then maintain, a level of force within a range indicated by a pair of blue force target rectangles. The robotic arm then automatically moved the cylinder to a target location (purple circle in Fig. 1, transporting) while maintaining the isometric force. When the arm reached the target location, the NHP released the robotic gripper by zeroing its isometric force output, placing the cylinder at the target location resulting in a successful trial. If the NHP completed the trial successfully, they received a juice reward based on the number of green cues displayed at the beginning of the trial, which were present throughout. If the animal failed the trial by not applying force at the proper time or by applying too much or too little force during the transport period, they received a timeout punishment based on the number of red cues, also initially displayed and visible throughout. After reward or punishment delivery, hereafter termed feedback, the robotic arm retreated automatically horizontally leftward to the starting position, and the subsequent trial began.

### Surgery

All animal experiments were performed per the ARRIVE guidelines. All surgical procedures were conducted in compliance with policies set forth by the National Institutes of Health Guide for the Care and Use of Laboratory Animals and were further approved by the State University of New York Downstate Institutional Animal Care and Use Committee. Once trained on the above task, two non-human primate (NHP) subjects (one male bonnet macaque (*Macaca radiata*) and one female rhesus macaque (*Macaca mulatta*)) were implanted in cS1, rM1, and PMd with 96-channel microelectrode arrays. These 10 × 10 Blackrock electrode arrays had electrodes spaced by 400 μm with lengths of 1.5 mm (rM1, PMd), or 1.0mm (S1), and 400 kOhm impedance with ICS-96 connectors. Implantation was conducted as detailed in our previous work (Chhatbar et al., 2010).

Briefly, NHP preparation and anesthesia were performed directly by the State University of New York Downstate Division of Comparative Medicine veterinary staff members. The researchers conducted the surgery under their supervision. Ketamine was used to induce anesthesia, and isofluorane and fentanyl were used for maintenance. Aseptic conditions were maintained throughout surgery as a craniotomy window was created over the target location. A probing electrode was used to identify the hand and forearm region of cS1. An electrode array was implanted at this site and immediately across the central sulcus for the corresponding M1 region. The PMd array was placed just dorsal to the spur of the arcuate sulcus unless vasculature was limiting access, see Fig.S.1. Dexamethasone was used to prevent inflammation during the procedure, and diuretics such as mannitol and furosemide were available to reduce cerebral swelling further if needed. Both subjects were observed hourly for the first 12 hours after implantation and were provided with a course of antibiotics (Baytril and Bicilin) and analgesics (buprenorphine and Rimadyl).

### Extracellular unit recordings

After a two-to three-week recovery period, spiking and local field potential (LFP) activities were simultaneously recorded with a multichannel acquisition processor system (MAP, Plexon Inc.) while the subjects performed the experimental task. Neural signals were amplified, bandpass filtered from 170 Hz to 8 kHz, sampled at 40 kHz, and thresholded to identify single units. Single- and multi-units were sorted based on their waveforms using principal component (PC)-based methods in Sort-Client software (Plexon Inc.) followed by offline spike sorting (Plexon Inc.) to identify primarily putative single units.

### Divisive normalization modeling

Reward and punishment encoding models have the general form, *f*_*ij*_ = *g*(*r*_(*i*)_, *p*_(*i*)_), for the i^th^ trial and the j^th^ unit. *r*_(*i*)_ and *p*_(*i*)_ are the cued reward and punishment levels for the i^th^ trial that can take values of [0,1,2,3]. These models were fit to individual unit firing rate data from PMd, rM1, and cS1. *f*_*ij*_ is the post-cue or post-feedback firing rate for the j^th^ unit during the i^th^ trial, with reward level *r*_(*i*)_ and punishment level *p*_(*i*)_, and *g* was one of the functions described below. For post-cue analysis, the firing rate was averaged over the 500 ms window following cue presentation. In this period, the virtual robotic arm was stationary or autonomously moving horizontally to the right during the reaching scene, approaching the cylinder at a constant speed, and the NHPs were not yet applying isometric grip force. For post-feedback analysis, firing rates were averaged over 500 ms following the indication of the result (reward or punishment); thus, there were no temporal confounds due to the different reward and punishment delivery periods.

We designed one linear model (model 1) and two different divisive normalization models (model 2 and 3) for our data analysis below. We utilized four possible reward levels (*r* ∈ [0,1,2,3]) and punishment levels (*p* ∈ [0,1,2,3]). The total number of trials with a particular reward and punishment level combination is referred to as *T*_*rp*_. In our task, the number of trials for each *rp* combination was not always the same. Therefore, we used weighted least squares (WLS) to avoid overfitting some reward and punishment level combinations instead of ordinary least squares (OLS). OLS tries to minimize the sum squared error *err* = ∑_*i*_(*y*_*i*_ − *f*(*x*_*i*_))^2^. Here, *y*_*i*_ is the neural data, *f*(*x*_*i*_) = *ŷ*, which is the model estimate for data point i. WLS tries to minimize the weighted sum squared error, *err* = ∑_*i*_ *w*_*i*_(*y*_*i*_ − *f*(*x*_*i*_))^2^, where *w*_*i*_ is the weight for the i^th^ trial type. *w*_(*i*)_ was based on the total trial number for that reward and punishment level on the i^th^ trial, *r*_(*i*)_, *p*_(*i*)_. If *r*_(*i*)_ = 0 and *p*_(*i*)_ = 0, *w*_(0,0)_ was defined as 1. For all other combinations of *r*_(*i*)_ and *p*_(*i*)_, *w*_*r*(*i*)*p*(*i*)_ was equal to 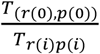. Thus, the total summed weighting for each reward and punishment level were all equal to 1.

**Model 1**: The first model assumed that a unit’s firing rate was the result of a linear relationship between reward and punishment,

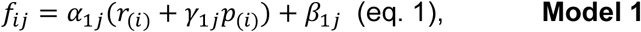

where *f*_*ij*_ was the post-cue or post-result firing rate for the j^th^ unit in the i^th^ trial. *r*_(*i*)_ and *p*_(*i*)_ are the reward and punishment levels, respectively, and *α*_1*j*_ and *γ*_1*j*_ are scaling factors with offset *β*_1*j*_. Model #1 is linear; however, note that the divisive models 2 and 3 can also show linear relationships between the stimuli *r*_(*i*)_ and *p*_(*i*)_ and the firing rate. There were four possible reward and punishment levels. 16 combinations were making up the affective stimuli and induced neural states. For each trial (i), the reward and punishment levels were chosen pseudo-randomly from a range of 0 to 3. Equation (1) was fit to firing rates from units for each region separately (cS1, rM1, and PMd). Adjusted R squared values were calculated for all models to determine which best explained the data (see below).

**Model 2**: The second model was a divisive normalization model incorporating reward and punishment levels,

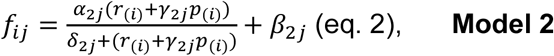

where *f*_*ij*_ was the post-cue, or post-result firing rate (500ms bin) for the j^th^ unit in the i^th^ trial, and *r*_(*i*)_ and *p*_(*i*)_ were the reward and punishment levels for that i^th^ trial, respectively. *γ*_2*j*_ was a scaling factor between reward and punishment for the j^th^ unit and *δ*_2*j*_ was an additive factor. Equation (2) was used to determine if the j^th^ unit encoded valence or motivation. If *γ*_2*j*_ < 0 (eq.2) the unit was considered to encode valence, where reward and punishment modulated the firing rate in opposite directions. If *γ*_2*j*_ > 0 the unit was thought to encode motivation, as both punishment and reward modulated in the same direction since both reward and punishment are motivating factors (see Fig.S.2 for statistical testing flow chart). Estimated values of *α*_2*j*_, *β*_2*j*_, *γ*_2*j*_, and *δ*_2*j*_ were determined with weighted nonlinear least-squares fit (Matlab curve fitting app, function “fitnlm”), where 30 random initializations were utilized and the best among the 30 chosen as the best fit model (see Fig.S.2).

We utilized trial duration as a behavioral measurement and proxy for the NHPs affective state. We fit the trial duration data to a modified equation (2) for trial duration (*t*_(*i*)_) for a given reward level *r*_(*i*)_ and punishment level *p*_(*i*)_ combination under the reasonable assumption that these sensorimotor brain regions are at least partially responsible for the behavioral output from the NHPs, and thus if the neural activity is normalized, so should the behavior.

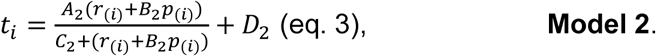

**Model 3**: For the third model, the denominator from equation (2) was modified to incorporate neural population information. We have

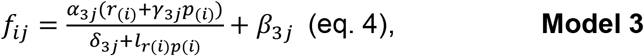

where *r*_(*i*)_ = [0, 1, 2, 3] and *p*_(*i*)_ = [0, 1, 2, 3] are the cued reward and punishment levels on the i^th^ trial. Like model 2, to avoid overfitting, we used WLS. The j^th^ unit’s post-cue or post-feedback firing rates with reward level *r*_(*i*)_ and punishment level *p*_(*i*)_ noted as *f*_*ij*_, were used to fit model 3. *l*_*r*(*i*)*p*(*i*)_ was a scalar based on the population firing rate for the combination of reward and punishment levels (*r*_(*i*)_, *p*_(*i*)_). *l*_*r*(*i*)*p*(*i*)_ was calculated using the following equation:

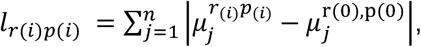

where j is the index of units and N is the total number of units. 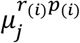 is the mean post cue or post feedback firing rate for the jth unit on trials with reward level *r*_(*i*)_ and punishment level *p*_(*i*)_, 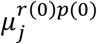 is the mean firing rate for the jth unit when *r* = *p* = 0. Intuitively, *l*_*r*(*i*)*p*(*i*)_ is a representation of the difference between the population mean firing rate when the reward level is *r*_(*i*)_ and punishment level is *p*_(*i*)_ as compared to the affective “baseline” *r* = *p* = 0. Using the raw mean firing rate was not as robust compared to the absolute value, as individual units may modulate in opposing directions in response to reward or punishment, thus canceling each other’s changes out. Estimated values of *α*_3*j*_, *β*_3*j*_, *γ*_3*j*_, and *δ*_3*j*_ were determined by fitting the model to the data as with model 2. We also tested the simple mean firing rates as normalization terms for comparison.

We compared fitting results using the adjusted *R*^2^ between mean firing rates and absolute differences. All significant units had higher adjusted *R*^2^ values using the absolute value difference as compared to the simple mean rate. We continue in this paper considering only the absolute value difference form of model 3, as part of our goal is to find the best model form that could be used in a brain-machine interface (Marsh et al., 2015; An et al., 2018; Zhao et al., 2018). Significant unit here means a unit had residuals with a normal distribution (JB test), passed an F-test, and had at least one parameter related to reward or punishment as significantly non-zero (please see next paragraph for more detail and Fig.S.2).

### Motivational intensity and valence encoding

Motivation here is defined as *γ*_3*j*_ > 0, *γ*_2*j*_ > 0 or *γ*_1*j*_ > 0 for a given unit, meaning that reward and punishment both modulated the firing rate in the same direction. Valence is defined as *γ*_3*j*_ < 0, *γ*_2*j*_ < 0 or *γ*_1*j*_ < 0 for a given unit, meaning that reward and punishment had the opposite modulation for a given unit. Notice that one unit can be classified as motivation for one model and valence for another (see Fig.6, M1 units). We wanted to pick the best model for each unit and make sure it was significant, and then determine if that model was for motivation or valance and report the model’s predictive adjusted R^2^ for comparisons.

To determine significant units the following was conducted for every unit: 1) We obtained fitting results using models 1, 2, and 3 starting each model at 30 random initial conditions. We utilized a 10-fold cross validation procedure for the best set of parameters for each of models 1, 2, and 3. 2) We picked the best model (the model with the highest 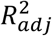, see eq (10) for model comparisons). 3) We conducted statistical tests for the best model. The statistical tests contained the three following parts: 1) We tested if the residuals for the best model had a normal distribution (Jarque–Bera test (JB), p<0.05). If this model successfully passed the first test we moved on. 2) We ran an F-test for that model to see if that model was significantly different from a constant model (p<0.05). The constant model was defined as *f*_*j*_ = *c*_*j*_, where *c*_*j*_ was the mean firing rate for all data points (without regard to r and p levels). We calculated the F value by:

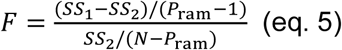

where *SS*_1_ was the weighted sum of the squared error from the constant model, *SS*_2_ was the weighted sum of the squared error from the fit model, *P*_*ram*_ was the number of parameters from the fit model, and *N* was the number of data points. If this model successfully passed the F test in the second step, we continued. Keep in mind that as we move forward, the model has sequentially passed X statistical tests at p<0.05. Therefore, the actual cumulative probability is a product of these tests applied sequentially, which we will term p_seq_ that equals 0.05 * 0.05 at step #2 or p < 0.0025. 3) We then tested if the parameters related to reward or punishment were significantly different from zero, and that the *γ* parameter was significant as indicated by the output of Matlab’s fitnlm function, 2 t-tests at p<0.05 and p_seq_ < 0.000006. If true, we defined this unit as significant and moved on. 4) We then found if that unit was only significant for reward encoding (only parameters for reward level were significantly different from zero), only significant for punishment encoding (only parameters for punishment level were significantly different from zero), or significant for both reward and punishment encoding. 5) For units that were significant for both reward and punishment encoding (p_seq_ < 0.0000003), we then determine if the unit was motivational or valence encoding using that best significant model’s significant *γ*. See Fig.S.2 for a graphical representation of these best-case probabilities assuming independence. For all significant units whose best model was model 2, additional analysis was conducted to determine if that unit was hyperbolic or not. A unit is hyperbolic when the fitted model 2 for the unit had a singularity, where the function goes to infinity, and that singularity fell inside our stimulus range (R from 0 to 3 and P from 0 to 3). However, it should be noted that our affective stimuli formed a discrete set, and we would not see an infinite output from the model at one of the actual affective stimuli. This is the case as we were minimizing a sum of squares, and clearly, even a single infinite output would never be a minimum in this case. So, these points where the result would diverge to infinity should not be taken too seriously.

Post-feedback data analysis was similar to the post-cue analysis above. Models 2 and 3 were fit to post-feedback data as with the post-cue period. Equations (2) and (4) were used, where *r*_(*i*)_ = [0,1,2,3] and *p*_(*i*)_ = [0,1,2,3], and *f*_*ij*_ were the post-feedback firing rate for the j^th^ unit at the ith trial with reward level *r*_(*i*)_ and punishment level *p*_(*i*)_. This feedback delivery period was the 500 ms window following the onset of reward or punishment delivery. For the zero-reward, zero-punishment trials, the point in the trial at which reward or punishment would have been delivered was used. Like the post-cue analysis, we picked the best model using the above-outlined methods where several questions are asked in order leading to highly significant p_seq_ < 0.0000003. A difference between the post-cue and post-feedback analysis was at the beginning of the trial (post-cue), the animals were likely to incorporate both reward and punishment expectations, as they did not know the outcome of the trial; thus, there was uncertainty. The post-feedback period was after the animal had successfully completed or failed the trial, and the animal at that point may only be encoding reward for successfully completed trials or punishment for unsuccessfully completed trials; at least, this was our hypothesis. Thus, we fit encoding models for successful trials and failed trials separately to test this hypothesis. For successful trials, the model was based on reward-only modulation for post-reward data:

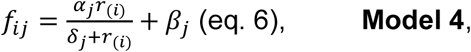

for successful trials only and for failed trials, the model was based on punishment-only modulation,

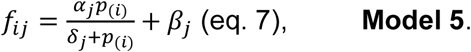

Where *r*_(*i*)_ or *p*_(*i*)_ were the reward or punishment level for the i^th^ trial, and *f*_*ij*_ was the post-feedback firing rate for the j^th^ unit in the i^th^ trial. *δ*_*j*_ and *β*_*j*_ were parameters fit with nonlinear least-squares fitting as before. Equation (6) was fit using all data from all successful trials. Equation (7) was fit using all data from all failure trials. Like models 2 and 3, we can design models 6 and 7 using information from population firing rates as the divisive terms instead of the current reward or punishment level. For successful trials, we have

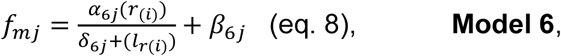

where 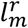 is the scalar based on reward trial population firing rates when, *l*_*r*(*i*)_ was the mean absolute value firing rate difference compared to the baseline, which was r = 0, similar to Model 3. Likewise, for failed trials, we have

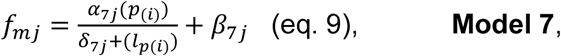

where again*l*_*p*(*i*)_ is the scalar based on punishment population firing rates when, *l*_*p*(*i*)_ is the mean absolute value firing rate difference compared to the baseline, p=0. WLS was used for models 4∼7 to avoid overfitting particular reward or punishment levels.

### Comparing fitting results across models

Models used did not always have the same numbers of parameters, therefore the adjusted 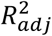 values (10-fold cross-validation) were calculated and used to compare the models. 10-fold cross-validation is a procedure where the original dataset is split into 10 equal partitions, and from these, 9 are used for fitting the model. The 10^th^ is used for determining how well the model generalizes to unseen data. This is then repeated 10 times such that every data partition is used once as the held-out data to determine the model’s generalizability. The 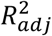 was determined by,

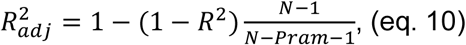

where *N* is the sample size for all models, and *Pram* is the total number of explanatory variables/parameters in the model, which was *n* = 2 for models 1, 4, 5, 6, and 7, and *n* = 3 for models 2 and 3. During the model comparison, we first picked the best model by finding the model with the highest 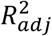 (see eq. 10). Then for this model, we ran an F-test (see eq. 5) to determine if that unit was significant or not.

## Results

Before describing the divisive normalization analysis on cued reward and punishment trials, we show qualitative results in Fig.2 with simple peri-cue and peri-feedback time histograms. Example single units from the three brain regions (PMd, M1, and S1) for the two NHPs are separated into units that had a qualitative motivational intensity relationship with cued reward and punishment (Fig.2 NHP S S1 post-cue) or a valence relationship that is the neural rate either increased or decreased in a simple manner as trial valence increased, such as R0P3 with the lowest valence and R3P0 with the highest (Fig.2 NHP P M1 pre-result). Where R stands for reward value and P for punishment value, likewise, the motivational intensity would be highest for R3P3 and lowest for R0P0 and is a symmetric function compared to the valence function that would be linear from R0P3 to R3P0, in either an increasing or decreasing manner. Note the distribution of response types in Fig.2.

**Figure. 2.**
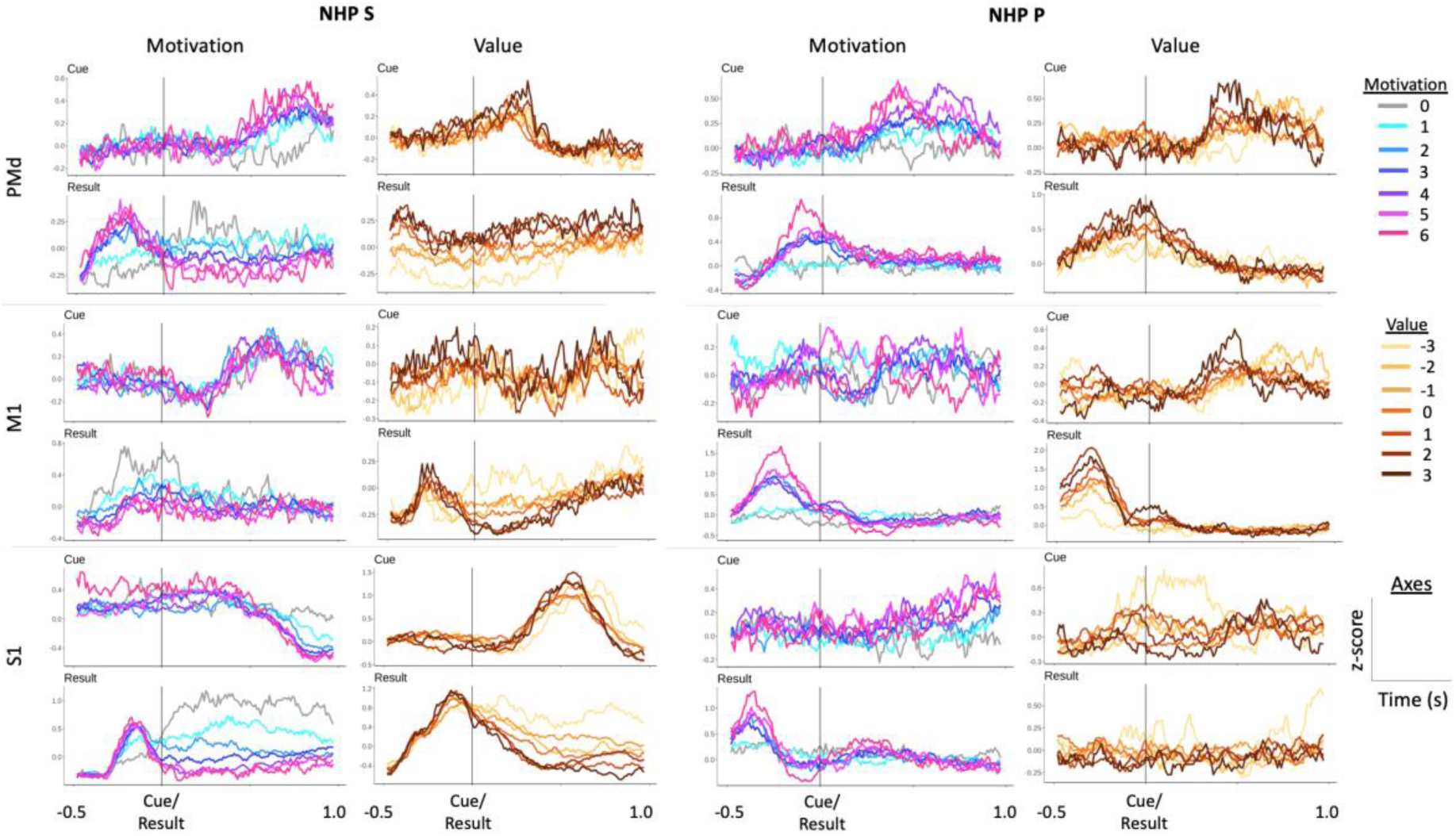
shows the peri-cue and peri-feedback time histograms for several example single units from each of the brain regions recorded from both NHPs. These units were chosen qualitatively. Each cue and result pair in a column are from an individual and distinct unit. Here we show units that qualitatively represent either valence-like or motivational-intensity-like response patterns. Units were binned and z-score normalized throughout a session. This normalized firing rate was then averaged for each valence level, defined as the reward level minus the punishment level for a given trial, or each motivational intensity level, defined as the reward level plus the punishment level for a given trial. Individual units demonstrated a range of responses following cue presentation and before and after feedback.

We recorded 3 blocks of data, from two different days, for each of the two NHPs. For each block, the number of R-level and P-level trials and the number of units for every region and NHP are shown in Tables 1 and 2, respectively. We tested several normalization models, one linear (model 1), and the rest divisive normalization models (DNM). As the divisive normalization models (DNMs) fit the data better ∼100% of the time, we focus on the DNMs. Below we first show results from a DNM that utilized the categorical R and P levels from the task (eq. 2), which was termed model 2 (Fig.3); second, results for model 3 (eq. 4) are shown in Fig.6. Model 3 utilized the population activity within a brain region in the divisive term (see methods). In Fig.3, we present the trial time results, using simultaneously cued R and P levels (model 2, eq.3). The rationale for fitting the trial times to the same DNM #2 was that we hypothesized the motor output from the sensorimotor regions are producing the isometric force output, at least in part. Therefore, the duration of a trial is dependent on the neural activity within these regions to an extent. Using the same model form, we could determine if the neural activity follows the same format as a behavioral measure governed at least in part by the neural activity within the regions under study. We found that trial durations always followed a motivational intensity form more clearly than a valence form. This behavioral measure indicates the NHPs understood the cues, as our task did not involve any choice other than the NHP deciding to either work or not.

**Table 1:** Trial number for each cued reward (R) and timeout punishment (P) Level

| Section 1 for NHP P |  |  |  |  |
| --- | --- | --- | --- | --- |
|  | R = 0 | R = 1 | R = 2 | R = 3 |
| P = 0 | 10 | 3 | 8 | 10 |
| P = 1 | 11 | 14 | 14 | 13 |
| P = 2 | 11 | 12 | 8 | 8 |
| P = 3 | 22 | 13 | 6 | 6 |
| Section 2 for NHP P |  |  |  |  |
|  | R = 0 | R = 1 | R = 2 | R = 3 |
| P = 0 | 41 | 5 | 16 | 7 |
| P = 1 | 22 | 8 | 13 | 6 |
| P = 2 | 7 | 5 | 11 | 5 |
| P = 3 | 3 | 10 | 11 | 3 |
| Section 3 for NHP P |  |  |  |  |
|  | R = 0 | R = 1 | R = 2 | R = 3 |
| P = 0 | 18 | 16 | 21 | 12 |
| P = 1 | 13 | 17 | 4 | 12 |
| P = 2 | 11 | 16 | 10 | 12 |
| P = 3 | 18 | 11 | 12 | 13 |

| Section 1 for NHP S |  |  |  |  |
| --- | --- | --- | --- | --- |
|  | R = 0 | R = 1 | R = 2 | R = 3 |
| P = 0 | 18 | 18 | 17 | 21 |
| P = 1 | 28 | 18 | 20 | 28 |
| P = 2 | 15 | 21 | 21 | 19 |
| P = 3 | 18 | 18 | 12 | 16 |
| Section 2 for NHP S |  |  |  |  |
|  | R = 0 | R = 1 | R = 2 | R = 3 |
| P = 0 | 12 | 17 | 12 | 17 |
| P = 1 | 14 | 21 | 14 | 17 |
| P = 2 | 13 | 15 | 13 | 14 |
| P = 3 | 15 | 13 | 15 | 21 |
| Section 3 for NHP S |  |  |  |  |
|  | R = 0 | R = 1 | R = 2 | R = 3 |
| P = 0 | 26 | 18 | 12 | 14 |
| P = 1 | 17 | 25 | 19 | 34 |
| P = 2 | 24 | 14 | 17 | 22 |
| P = 3 | 14 | 11 | 18 | 17 |

**Table 2:** Number of Units for each recorded region for the two NHPs

|  | PMd | M1 | S1 |
| --- | --- | --- | --- |
| NHP P Section 1 | 140 | 77 | 75 |
| NHP P Section 2 | 83 | 67 | 78 |
| NHP P Section 3 | 85 | 77 | 91 |
| NHP S Section 1 | 85 | 57 | 81 |
| NHP S Section 2 | 87 | 58 | 85 |
| NHP S Section 3 | 109 | 72 | 93 |

In Fig.3.1, we see that NHP P was very sensitive to differences between rewarding (RX) and non-rewarding (R0PX) trials, where X is a placeholder for any level (0-3). In addition, there was an influence on trial time due to the cued punishment level Fig.3.1, which was most apparent when the reward level was zero. In these Fig.3 plots, the x-axis is termed the affective stimulus and combines the cued reward level and a scaled version of the cued punishment level. This scaling was necessary as the punishment was in timeout periods while the reward levels were in juice volume. In addition, reward and punishment are known to be represented by differing neural circuits. NHP S had a smoother relationship between affective stimulus and trial time. These results demonstrate that the mean trial time can be explained with a motivational intensity model, as *B*_2_ > 0 for both NHPs in equation 3. Thus, reward and punishment levels were added together, as expected for motivational intensity, compared to valence, where reward and punishment levels would be subtracted. The mean trial time decreases as the motivational intensity increases. In Fig.3.2, we plotted the neural firing rates vs. the affective stimulus, derived from the data for a given model that determined the relative scaling of punishment compared to reward. Red squares ± the SEM for each category, of which there were 16, are plotted. In blue, we have plotted the fit to the data from model 2,

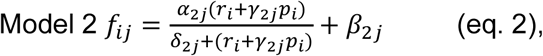

with fits that fall into one of three categories: First, essentially linear (Fig. 3.2 row 1), second, sigmoidal (row 2), and third hyperbolic (row 3). These were seen in both animals and all three cortical regions investigated (PMd, M1, and S1). Note that the aforementioned “linear response” is from the above DNM #2 that is plotted against the affective stimulus (see Fig.3.2 x-axis) and should not be confused with the simple linear relationship between a unit’s rate and the stimulus as described by model #1 in the methods section. In over 99% of all units, model 1 was inferior to the DNMs, and thus we do not focus on model 1, see table 3 for best model %.

**Table 3.** Percentage of all recorded units with better adjusted R-squared for given model comparison. PC = post cue, PF = post feedback.

| NHP P |  |  |  |  |  |  | NHP S |  |  |  |  |  |  |
| --- | --- | --- | --- | --- | --- | --- | --- | --- | --- | --- | --- | --- | --- |
|  | PC PMd | PC M1 | PC S1 | PF PMd | PF M1 | PF S1 |  | PC PMd | PC M1 | PC S1 | PF PMd | PF M1 | PF S1 |
| Model 2 better than 1 | 100% | 100% | 100% | 100% | 100% | 100% | Model 2 better than 1 | 100% | 100% | 100% | 99% | 100% | 100% |
| Model 3 better than 1 | 100% | 100% | 100% | 100% | 100% | 100% | Model 3 better than 1 | 99% | 100% | 100% | 99% | 100% | 100% |
| Model 3 better than 2 | 53% | 66% | 87% | 38% | 62% | 67% | Model 3 better than 2 | 49% | 64% | 61% | 41% | 35% | 49% |

**Figure 3.**
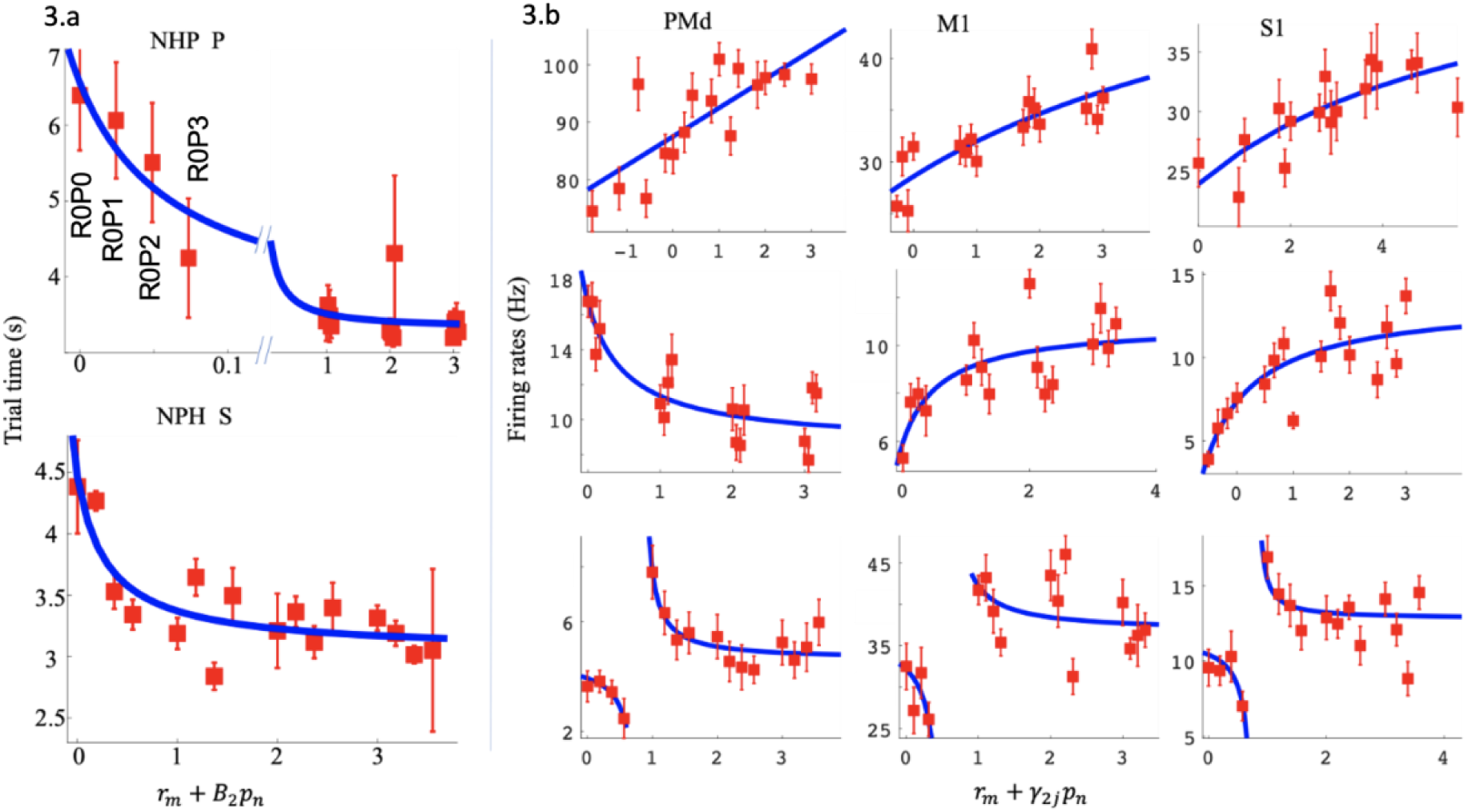
Post-cue reward (R) and punishment (P) analysis with divisive normalization model 2. Figure 3.a depicts the average and SEM of the trial time (red) for different affective stimuli. The x-axis represents the R and P level in the form of an affective stimulus *r*_*m*_ + *B*_2_*p*_*n*_, and the y-axis represents the trial time (s). We have labeled the R and P levels for the R = 0 group. Figure 3.b shows example unit R and P modulation for post-cue neural spike data. Each unit has an R and P encoding model 2 significantly different from a constant model (F-test, p<0.05).

For each subplot, the x-axis represents the affective stimulus as above, and the y-axis represents the post-cue firing rate (Hz). Each red point represents the mean post-cue firing rate (0-500ms post cue onset) for that R and P level ± the SEM, and the blue line represents model 2 fit to that unit’s data. The first column includes units from PMd, the second column M1, and the third column S1. The rows from top to bottom show examples of linear, sigmoidal, and hyperbolic units.

### Comparing linear (1) and divisive (2-3) models of valence and motivation

We picked the best model from models 1 (linear, eq.1), 2, and 3 (divisive, eqs.2, 4) for each unit. There were many units whose best model was model 2 (see table 3 and Fig.S.5). For these units, we analyzed if they had hyperbolic relations to affective stimuli or not. The results are shown in Fig.4. Overall, 36% - 100% of the units for a given brain region were hyperbolic with an average of 63% of all units being hyperbolic for model #2. Thus, there was a large portion of nonlinear modulation for units whose best model was model 2, with PMd having a clear majority of hyperbolic units in both NHPs post-cue with 100% in NHP P during the post cue period.

**Figure. 4.**
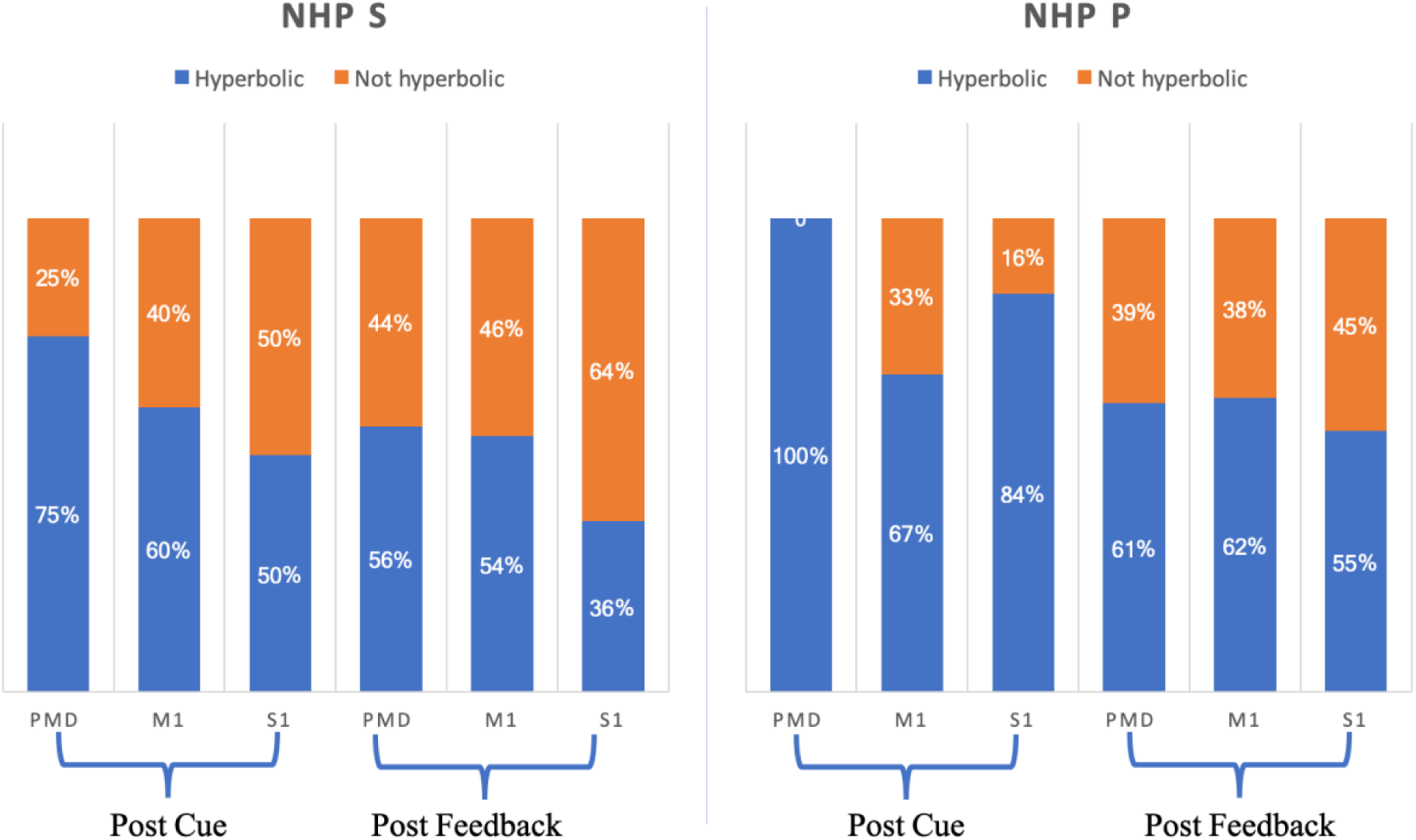
Number of significant units from model 2 that were hyperbolic.

For non-hyperbolic units seen in Fig.4.red, we asked how many units had a sigmoidal vs. approximately linear relation between affective stimuli and firing rate? To classify the distribution of these curve shapes as linear and sigmoidal units, we first found all significant units whose best model was model 2, 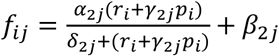. We analyzed all of the non-hyperbolic unit curve shapes by comparing the two slopes at the start and endpoint of the curve, as seen in examples (Fig.5). We then took the angle of the tangents at A and B, noted as *θ*_*A*_, *θ*_*B*_ respectively. We calculated the angle difference Δ*θ* = |*θ*_*A*_ − *θ*_*B*_| and used this as a proxy for the curves’ shape, linear or sigmodal. If Δ*θ* was “large” (> 30°), the shape was arbitrarily considered sigmoidal, based on the curves seen in Fig.5.a. If Δ*θ* was < 30° to zero, the shape of the curve was deemed linear. We combined all data for each NHP to have enough data points for a distribution of the Δ*θs*. For NHP S, about 25% of the non-hyperbolic units had a Δ*θ* larger than 30°. For monkey P, about 37% of the non-hyperbolic units had Δ*θ* larger than 30° as seen below in Fig.5. Thus, adding the hyperbolic and the sigmoidal unit groups to determine the number of nonlinear responses led to most units being classified as nonlinear in the affective stimulus vs. unit rate space.

**Figure. 5.**
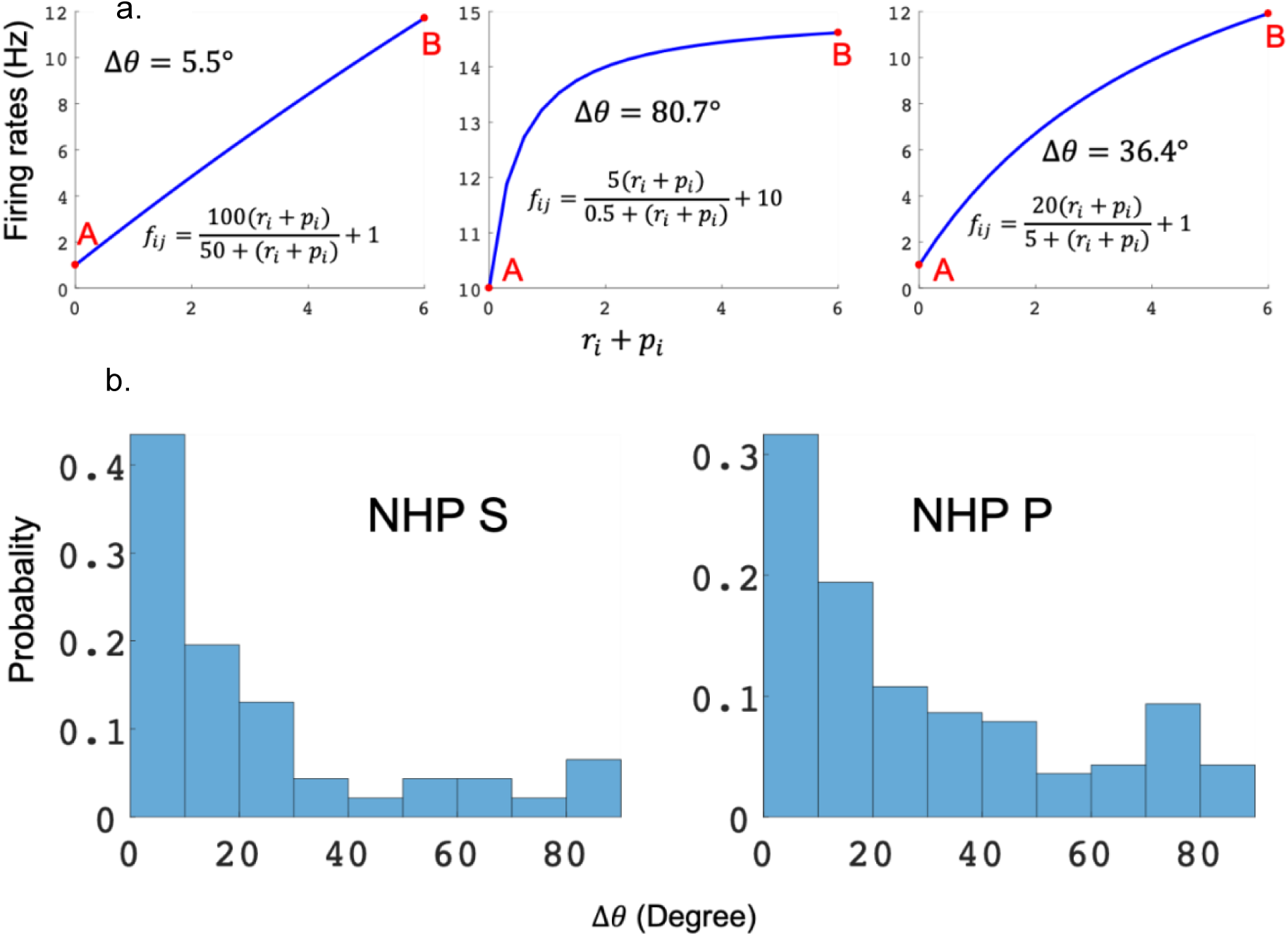
(a.) Description of method used to determine % of the population with sigmodal or linear response curves from the population that were not hyperbolic. (b.) Distribution of Δ*θ* for the non-hyperbolic units using model 2. Note that most non-hyperbolic units had a Δ*θ* < 30° making their responses approximately linear. All units shown here were not hyperbolic, that is, the red portion of the bars seen in Fig. 4 above.

As model 2 utilized the categorical R and P levels in the denominator, which act as the divisive term, we wished to determine if a more “natural” divisive term could perform better. We derived the divisive term for model 3 from the given cortical regions population activity as described in the Methods section; Model 3 is 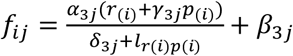 (eq. 4). In Fig.6, we show a comparison, for example units, between models 2 and 3. Model 3 had a higher fitting accuracy for all example units shown and visually better “explained” the modulation due to the affective stimuli. Units could have two general characteristics to their responses: focal tuning to a small reward and punishment range, termed hyperbolic, or a more gradual response to a broader range of reward and punishment levels, sigmodal or linear. Model 2 could only capture the focal tuning (hyperbolic) or the broader modulation (linear or sigmoidal units). However, model 3 could capture both modulation types, as seen in Fig.6 (see units in green circles). Here we show the qualitative results for a set of examples and follow with the population results and the model comparisons. Note the x-axis, which represents the affective stimulus, is fit to the data and can change between models 2 and 3, as can the unit’s apparent representation, valence vs. motivation.

**Figure 6.**
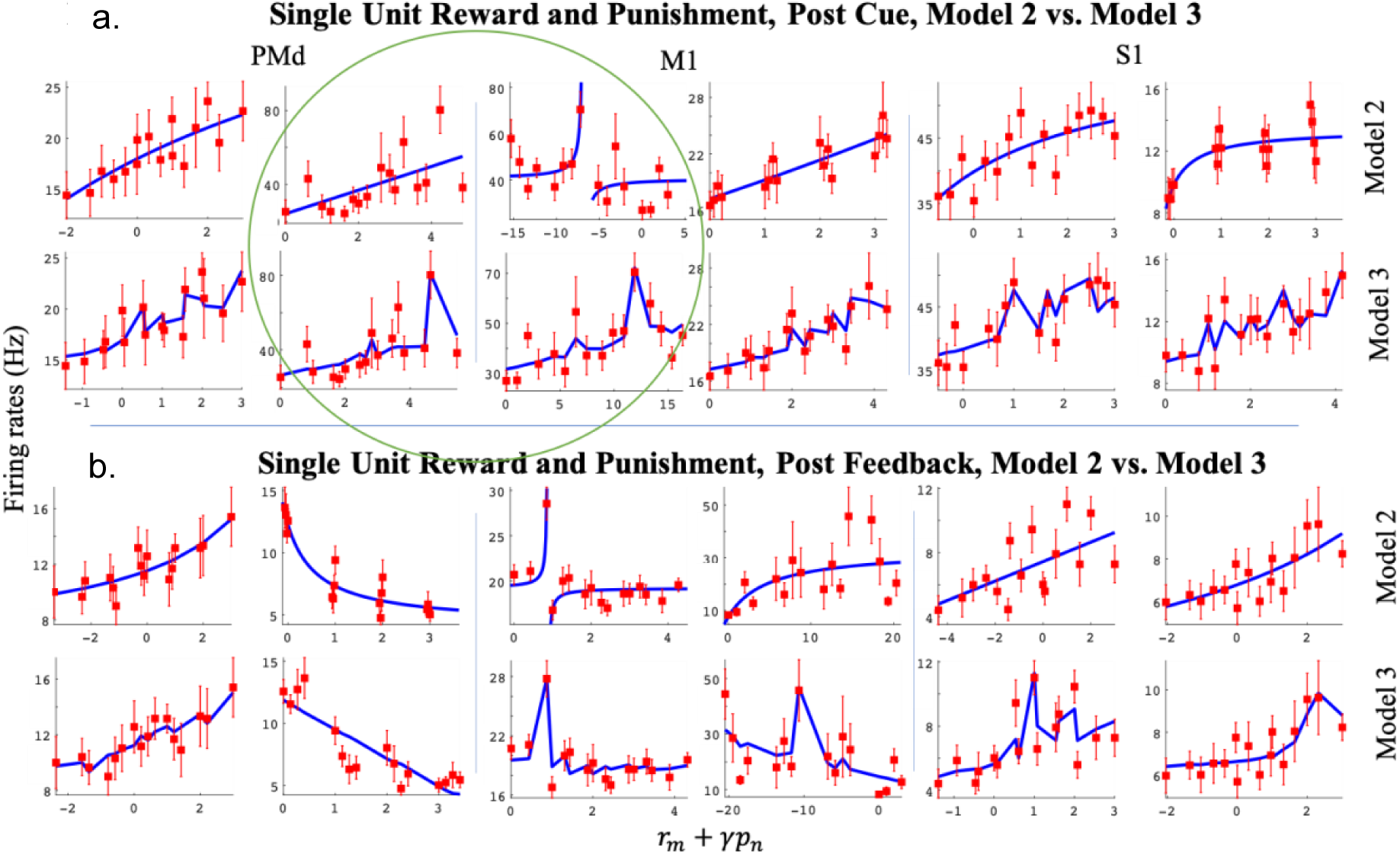
Post-cue (a) and post-feedback (b) example units for model 2 (top rows in each subplot) and model 3 (bottom rows in each subplot). The x-axis represents the affective stimuli, a linear combination of reward and scaled punishment *r*_*m*_ + *γp*_*n*_, where *γ* is fit to the data. The y-axis represents the post-cue (a.) and post-feedback (b.) firing rate (Hz). The mean ± SEM of the firing rates (red) and the mean for the model (blue) are shown. Each column represents one unit, fit to model 2 (top) and model 3 (bottom) rows. The first two columns show units from PMd, the third and fourth columns from M1, and the last two columns from S1. All units shown here were significant (see methods).

For each unit, we first found the best model among models 1, 2, and 3, where model 1 is a linear non-divisive form, Model 1 is *f*_*ij*_ = *α*_1*j*_(*r*_*i*_ + *γ*_1*j*_*p*_*i*_) + *β*_1*j*_ (eq.1). Second, we tested for each best model if the parameters related to reward and punishment were significantly different from zero, such that their confidence intervals for the given parameter did not overlap zero (see methods and Fig.S.2). This led to four cases: 1, parameters related to reward and punishment were both not significant. 2, only the parameter related to reward was significant. 3, only the parameter related to punishment was significant. 4, Parameters related to reward and punishment were both significant. Group 2, 3, and 4 were considered significant units. Only units from group 4 (red pie charts in Fig.7.b) could be motivation or valence encoding. Overall, we found approximately 25% - 48% of the units from the full population of a given brain region were significantly modulated by reward, punishment, or both. For NHP S, we had 127 significant units among 283 total units in PMd (45%), 63/187 units in M1 (34%), and 125/259 units in S1 (48%). For NHP P, we had 106 significant units among 308 total units in PMd (34%), 67/221 units in M1(30%), and 61/244 units in S1 (25%).

**Figure 7.**
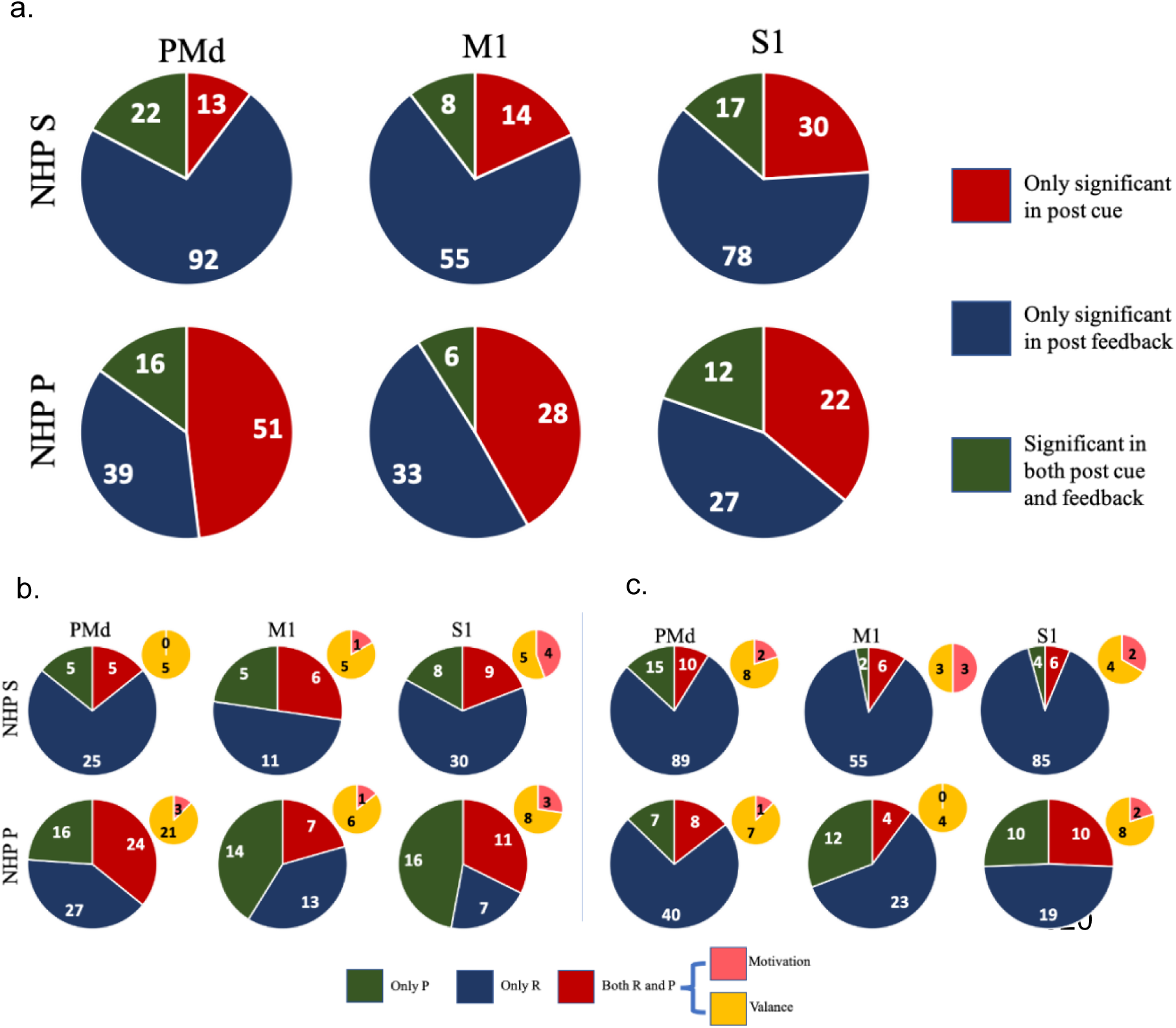
(a.) Task period distribution of significant units from two NHPs for three sensorimotor brain regions (PMd, M1, S1) fit to each unit’s best model. (b.) Post-cue reward and punishment encoding distributions of significant units using the units’ best model (see methods and text). (c.) Same as (b.), for the post-feedback period.

To determine if and how units changed their representation of reward and punishment throughout a trial, we looked at the units’ responses in the post-cue and post-feedback periods and plotted this information in Fig.7. Here we have broken the population of all units into the 4 groups mentioned above. We then further broke the units in group 4 down into motivational intensity and valence. From Fig.7, we can see that, in general, units modulate more for reward compared with punishment during the post feedback period. It also appears that more units represent valence in these data sets as compared to motivation. However, the number of units that we could use for this analysis was small. The units had to pass significance tests for multiple parameters simultaneously compared to the other categories. Nonetheless, we still obtained 106 total units from 3 brain regions in 2 NHPs that passed such significance criterion, of which 84, or 80%, showed a significant valence representation compared to 20% that showed motivational intensity. As seen in table 4, NHP P experienced more failed trials than NHP S, which could have led to NHP P showing a larger percentage of units with punishment-only representations, as seen in Fig.7.b.

**Table 4:** Trial numbers for each R and P level

| Successful trial numbers |  |  |  |  |
| --- | --- | --- | --- | --- |
|  | R = 0 | R = 1 | R = 2 | R = 3 |
| NHP P Section 1 | 42 | 36 | 25 | 25 |
| NHP P Section 2 | 30 | 31 | 36 | 26 |
| NHP P Section 3 | 33 | 45 | 41 | 39 |
| NHP S Section 1 | 70 | 74 | 67 | 83 |
| NHP S Section 2 | 58 | 62 | 54 | 59 |
| NHP S Section 3 | 77 | 63 | 57 | 77 |

| Failure trial numbers |  |  |  |  |
| --- | --- | --- | --- | --- |
|  | P = 0 | P = 1 | P = 2 | P = 3 |
| NHP P Section 1 | 10 | 13 | 8 | 10 |
| NHP P Section 2 | 10 | 28 | 8 | 4 |
| NHP P Section 3 | 19 | 14 | 10 | 15 |
| NHP S Section 1 | 9 | 5 | 3 | 4 |
| NHP S Section 2 | 6 | 7 | 4 | 4 |
| NHP S Section 3 | 7 | 10 | 6 | 6 |

We wished to determine if units up modulate or down modulate their neural activity for the different levels of reward, as well as punishment. A unit was significant only if the parameters related to reward or punishment were significantly different from zero (see methods). Thus, for a unit with a significant reward parameter, we tested if that unit up modulated for reward (firing rates for that unit increased when the reward level increased) or down modulated (firing rates decreased when reward level increased), similarly for the significant punishment units. The results are shown in Fig.8.a for reward analysis and 8.b for punishment analysis. Notice that the fraction of upmodulated units for reward modulation in Fig.8.a generally decreases from post-cue to post-feedback. For punishment, the trends are reversed with more units up modulated during post-feedback compared to the post-cue period. These trends are seen for both NHPs and most cortical regions.

**Figure. 8.**
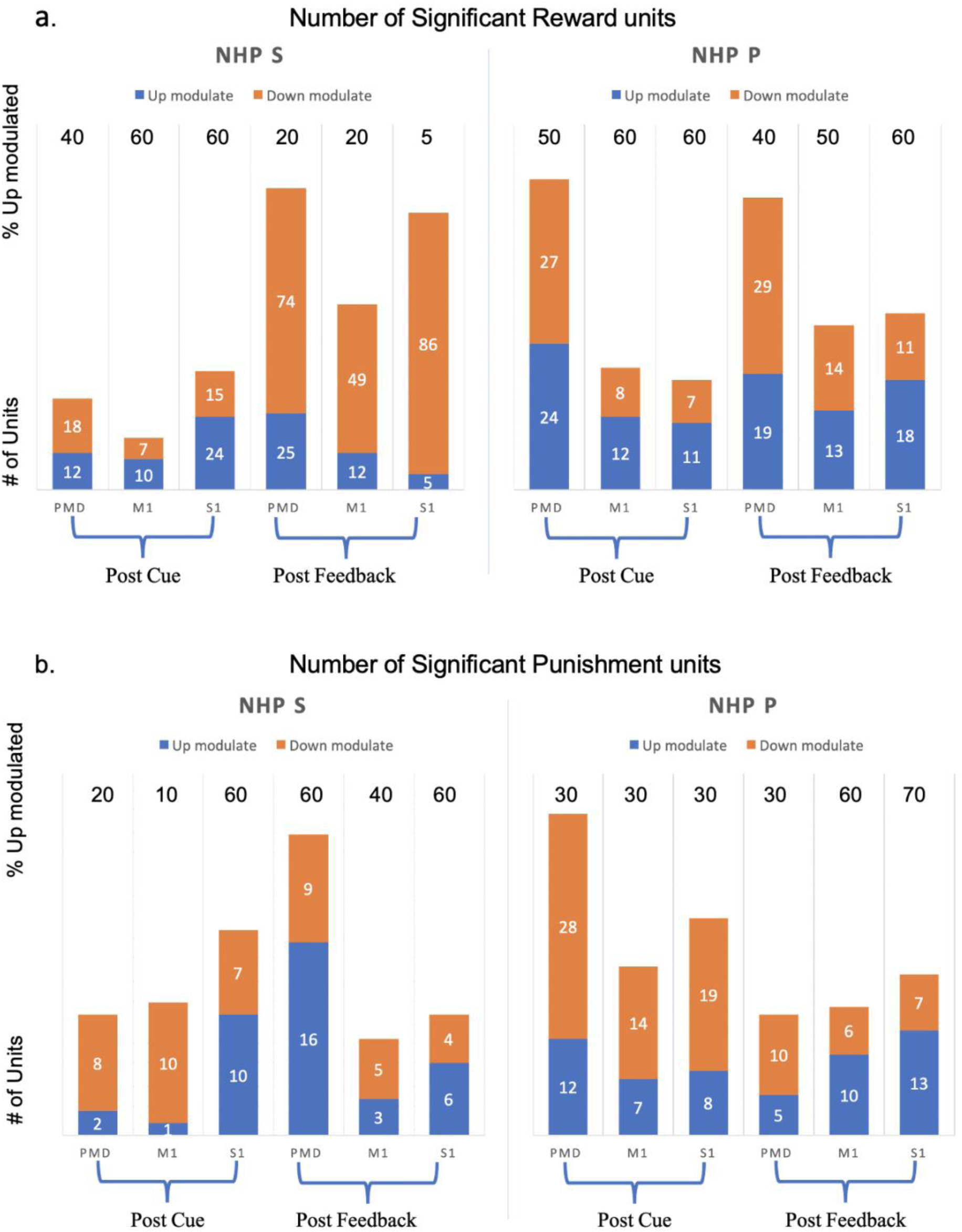
Modulation (up or down) for units with significant divisive normalization from two NHPs for three sensorimotor brain regions. (a.) shows population analysis for reward modulation. (b.) shows punishment modulation. Orange bars indicate the number of units with down-modulation, while the blue bars show the number of units with up modulation. Numbers at the top of each column are the percentage of units with up modulation.

Adjusted R-squared values for every unit using models 1, 2, and 3 for all regions have been compared. In all regions and periods, in essence, all units have better fits for the DNMs 2 and 3 as compared to the linear model 1. All data are shown in table 3 for both NHPs and the three brain regions (PMd, M1, and S1).

Furthermore, we conducted statistical tests to determine if two distributions (Fig.9) of 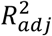 were significantly different from each other using the Wilcoxon Rank Sum test (p < 0.05). In general, it appears that NHP S’s data was more responsive during the post feedback period while NHP P was more responsive to the post cue period. NHP S’s results are: Median 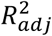 from PMd during post feedback was significantly higher than all 3 regions during the post cue period (pair-wise Wilcoxon Rank Sum test p < 0.05). The median 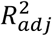 from M1, post feedback was significantly higher than all 3 regions during the post cue period. The median 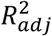 from S1, post feedback was significantly higher than S1 post cue. NHP P’s results are: Median 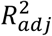 from PMd post cue was significantly higher than S1, post feedback. M1, post cue was significantly higher than PMd, post cue, and M1/S1, post feedback. S1 post cue was significantly higher than all 5 other datasets. PMd’s post-feedback median 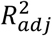 was significantly higher than S1’s post-feedback. Medians and ranges for all 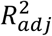 distributions are shown in figure 10.a. The corresponding proportion of significant units for all brain regions are shown in figure 10.b.

**Figure. 9.**
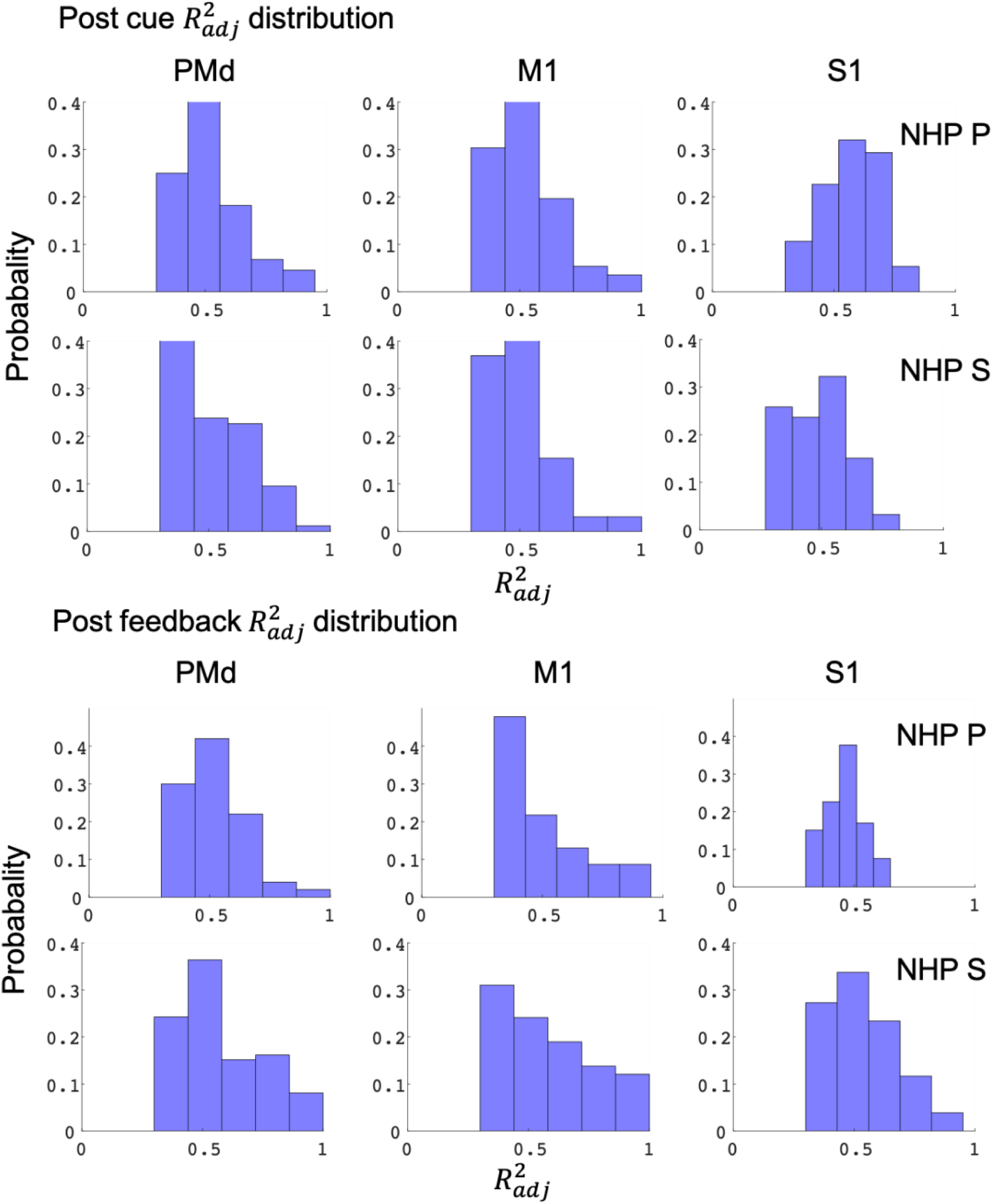
Distribution of the adjusted R squared values for the best significant model (model 2 and 3) fit for a given unit during the post-cue, top two rows, and the post-feedback, bottom two rows, for the two NHPs labeled on the right-hand side of the figure. Here we have fit the models to the mean firing rates for all 16 (R, P) levels; see supplementary figure S.Fig.4 for the fit distributions utilizing every trial rather than the 16 (R, P) categorical means.

**Figure. 10.**
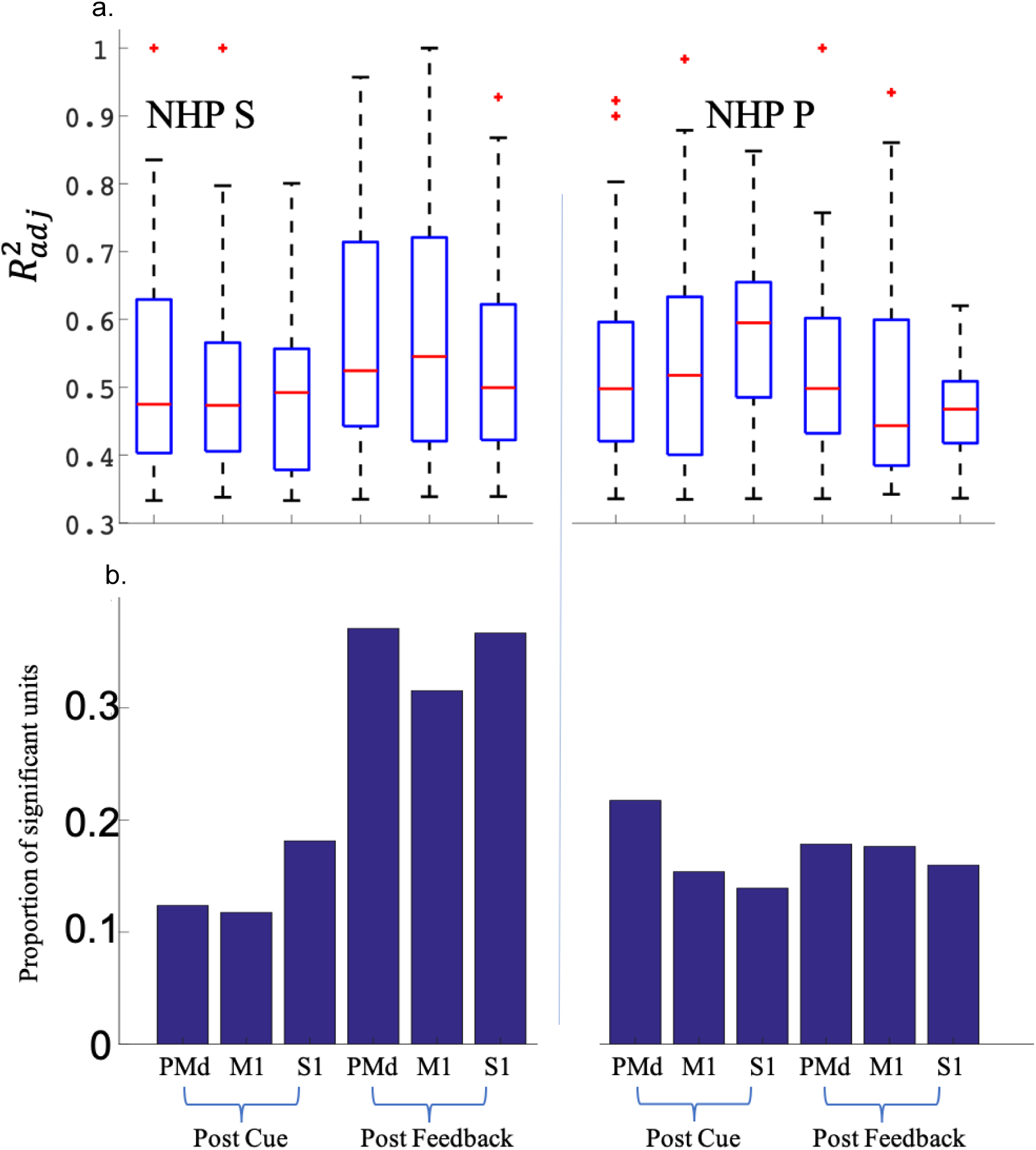
Medians and ranges for the adjusted R squared distributions are shown as boxplots (a.) see main text for statistical differences between regions and task periods. The proportion of significant units for all brain regions is plotted in (b.).

### Comparing divisive models (2-7) for reward and punishment feedback periods

Figure 7.b showed results for example units utilizing model 2 or 3 in the post-feedback window. For model 2 results, we plotted examples of sigmoidal and hyperbolic units. Post-feedback responses could be different from the post-cue responses for a given unit. Our hypothesis for this post feedback period was that the units would be encoding the reward or punishment that the NHP would receive compared to during the post cue period when the outcome was not yet known, and thus there was still uncertainty. Therefore, as an example, on successful trials in the post feedback period, we expected a unit to have the same response when the reward and punishment levels were R = 1, P = 0 or R = 1, P = 1 since the NHP would have obtained the same reward in either case. Therefore, if the neural activity represented valence, it should not change under these two example conditions, assuming limited history dependence. To test this assumption, we studied 4 different models for the post feedback analysis:

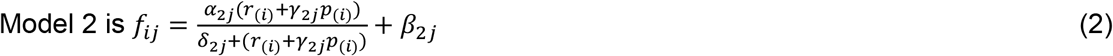

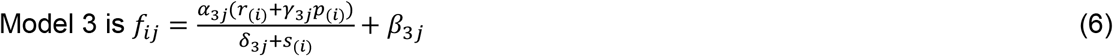

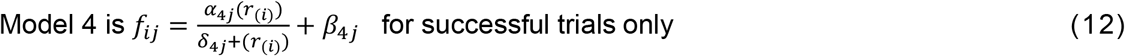

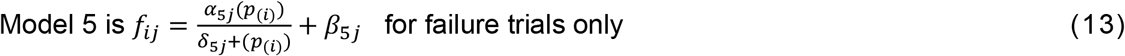

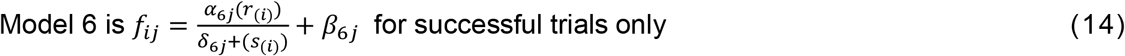

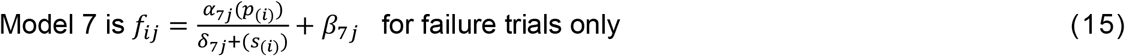

The adjusted R-squared was utilized for cross-modal comparisons. The results are shown in figure 11 and table 5. Overall, NHP P had slightly more units with their best model from group 1 (considering reward and punishment together for all trials). In contrast, NHP S showed the response we had anticipated, which is better fits in the post outcome period with group 2 (only considering reward for successful trials and only considering punishment for failed trials). Note that even though the two NHPs have slightly different patterns of responses, they are internally consistent between their brain regions, and these differences line up with the differences seen during unsuccessful trials. NHP S had 6% failures (out of 872 trials), while NHP P saw 27% (out of 558 trials). NHP S seemed to care more about the reward it would receive at the end of the task, perhaps because it could finish most trials successfully. NHP P had a more balanced exposure to successful and unsuccessful trials and had more units with their best fit were models 2 and 3 that included both types of trials (see table 5 for models 2-4, and Fig.11).

**Figure 11.**
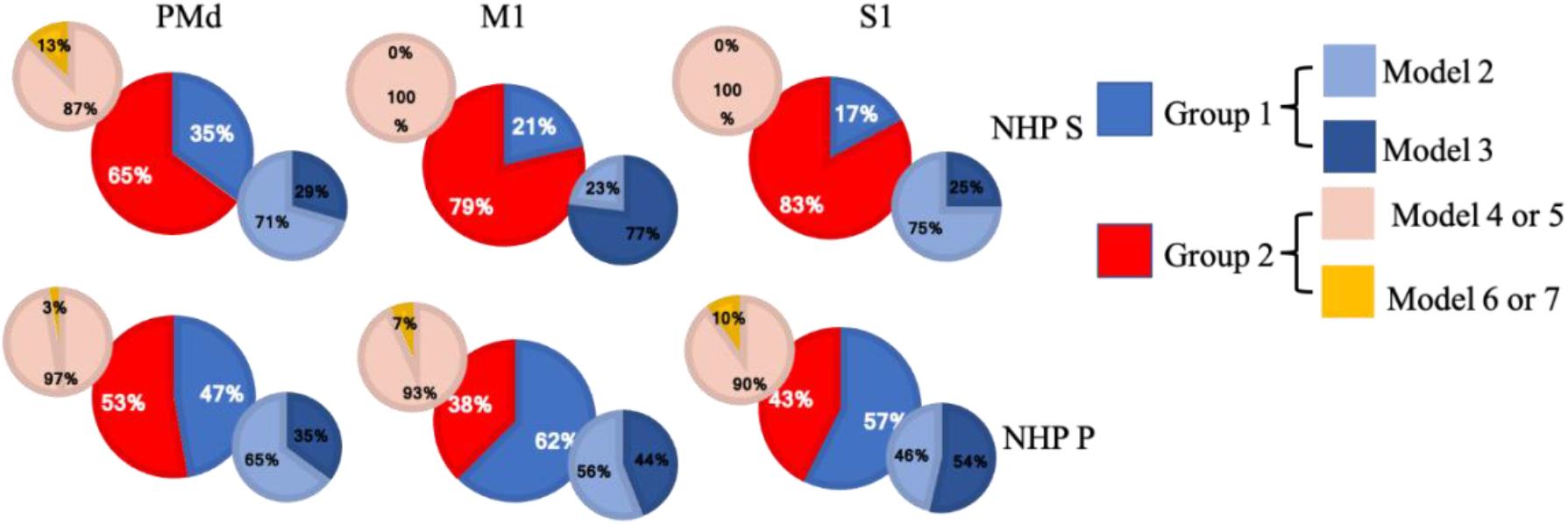
Percentage of units firing rate best fit for a given divisive normalization model. Group 1 models include both successful and unsuccessful trials, where model 2 utilized task R and P levels and model 3 utilized the population activity. Group 2 models include successful trials only (models 4, 6), or unsuccessful trials only (models 5, 7), with models 4 and 6 utilizing task R or P levels respectively, and models 5 and 7 using population activity in the divisive term from R or P trials respectively.

**Table 5:** Number of significant units with best model # (2-7), all were better than the linear model 1.

|  | Model 2 | Model 3 | Model 4 | Model 5 | Model 6 | Model 7 |
| --- | --- | --- | --- | --- | --- | --- |
| NHP S PMd | 24 | 10 | 48 | 7 | 2 | 6 |
| NHP S M1 | 10 | 3 | 48 | 0 | 0 | 0 |
| NHP S S1 | 12 | 4 | 74 | 3 | 0 | 0 |
| NHP P PMd | 22 | 12 | 37 | 0 | 1 | 0 |
| NHP P M1 | 14 | 11 | 11 | 3 | 1 | 0 |
| NHP P S1 | 13 | 15 | 16 | 2 | 2 | 0 |

## Discussion

Here we report widespread neural modulation of the sensorimotor cortices (cM1, rS1, and PMd) by simultaneously manipulating cued possible reward and punishment. Specifically, we found strong evidence for divisive normalization by valence and motivational intensity within these cortices. Two NHP subjects made cued isometric grip force “movements”. At the same time, we modulated the level of cued valence, spanning from negative (punishment) to positive (reward), and cued motivational intensity, which was simultaneously modulated by cueing the level of reward they would receive if successful, and level of timeout punishment if unsuccessful. There were 4 levels of cued reward and cued punishment, leading to 16 levels of valence and motivational intensity termed affective stimuli. We simultaneously recorded from 96 electrodes in each M1, S1, and PMd for 288 electrodes per NHP, a total of 576 electrodes.

We found three prominent relationships between neural firing rates and affective stimuli, linear, sigmoidal, or hyperbolic. Sigmoidal units had a two-part firing rate response, one high sensitivity, and one lower sensitivity region. Sigmoidal and linear units allow affect to modulate the neural rate over a wide range of stimuli, and the linear units may become sigmoidal if the affective stimuli have a broader scope. We suspect a distribution of units, some having wide affective modulation ranges, while others have more focal fields, such as the sigmodal and especially the hyperbolic units. Hyperbolic units were susceptible to a small range of affective stimuli, where the denominator in model 2 was close to zero. We could not determine the asymptotes for models 3, 6, and 7 as they utilized the population’s activity in the divisive term, unlike model 2 that used the R and P task levels. For model 2 we could interpolate to find the asymptotes. Model 2 may seem less biomimetic than models 3, 6 and 7; however, as we do not know precisely where the divisive information is determined, model 2 allowed us to ask questions about such normalization without assuming the local neural activity is performing this normalization. In this sense, model 2 is more powerful as it makes fewer assumptions.

In general, the population of units showed an increase in activity during the post-cue period and a significant suppression during the post-feedback period. During both the post-cue and post-feedback periods, we found the DNM (models 2-3) outperformed the linear model (model 1), as seen in table 3. This suggests that reward and punishment modulation can be nonlinear for PMd, M1, and S1, and DNM can capture this nonlinearity as well as linear relationships. Model 3 performed better than model 2 for several regions and times in NHP P (see table 3), suggesting that the accuracy of DNM incorporating affect can be improved by including population firing rates in the normalization term. Divisive normalization, scaled to the response of a population of units, has also been described in the invertebrate olfactory system and the primary visual cortex (Carandini & Heeger 2012). Thus, further supporting the idea that divisive normalization is a computational mechanism performed widely in various cortices. Also, of importance is that this type of normalization by the population can be more easily utilized toward stabilizing brain-computer interfaces in the face of changing affect, as discussed below (Zhao et al., 2018). However, more significant units had model 2 as their best model, which used the task labels to define the affective stimuli, compared to model 3, when including both NHPs, as seen in table 5. Model 4 had the most significant units as their best model, where model 4 only looked at successful trials and reward levels (R = 0-3) while also utilizing the task R levels in the divisive term.

### Valence and motivation

Previously, work on the influence of value in the sensorimotor stream has demonstrated value-based divisive normalization in the lateral intraparietal cortex (Louie, Grattan, and Glimcher 2011). This activity was interpreted as encoding the action value within the context of choices between actions associated with different state values. Later work argued against this interpretation in favor of motivational salience (Leathers and Olson, 2012). Subsequent work suggested the action value neural interpretation holds for behavior in general, including human subjects as well as NHPs (Louie et al., 2013). However, (Gluth et al., 2020) found that the behavioral responses were better fit with value-based attention when increasing the number of participants used in the previous study (Louie et al., 2013). However, see (Webb et al., 2020) for a contrary response to Gluth et al. in support of the divisive normalization point of view. We have shown in the current paper that we see both a valence-like signal and a motivational intensity-like signal. However, further work is needed to determine the exact non-movement-related variables encoded within these sensorimotor cortices. In one influential latent leviable model of affective space, two latent dimensions, by which all affective states can be represented, are arousal and valence (Russell, 1980). These two latent dimensions could be what we saw in our data.

In our study, the subjects did not have options to choose from and simply had one type of movement that they could perform successfully or not. Future work where we explicitly include choice should be helpful in further determining if both state-value/motivation and action-value/motivation are being represented in PMd, M1, and S1. Roesch and Olson studied prefrontal and frontal cortex activity in a set of papers, including premotor and supplementary motor regions. They found the increasing activity of these brain regions in line with increased motivational intensity as one moved more caudal (Roesch and Olson 2003) when comparing a small and large reward context. However, as only reward was modulated, they could not determine if it was valence encoding or motivational intensity. In later work, they included possible reward and punishment, the paradigm that inspired ours, and found results in the premotor region best fit by motivational intensity (Roesch and Olson 2007, 2004). However, this could partially be due to the analysis methods used or task-specific aspects. Their task included a memory component and choice.

In contrast, we focused on the influence that affective stimuli had on neural activity during a simple isometric grip force task. By utilizing divisive normalization models, we see a more robust valence representation while still seeing evidence of motivational intensity. Therefore, we claim that both valence and motivational intensity are being represented in PMd, M1, and S1 during our grip force task. In our data, valence was more strongly represented, see Fig.7.b. However, while looking at the larger picture from others’ work, task constraints may be influencing the outcomes, and thus we aim to conduct a comprehensive study in the future, including both tasks with and without choice. In addition, the range of the reward and punishment space should be modulated in blocks to determine how the contextual boundaries of R and P change the neural relationship to individual R and P levels. As we have suggested, this could lead to our linear units becoming sigmodal or even hyperbolic. This would allow us to ask questions about the distribution of the hyperbolic units over different affective spaces.

### Similarities between NHPs and brain regions

Fig.4 showed that both NHPs’ PMd had a high percentage of hyperbolic units using model #2, utilizing the task’s true R and P values. This is seen as the high percentage of significant hyperbolic units (75% NHP S and 100% NHP P) during the post reinforcement cue period. In general, most regions in both NHPs had more than 50% of their units as hyperbolic, which means the units were activated mainly by a given R and P level. Among the non-hyperbolic units, red portion of bars in Fig.4, the majority had a linear response to affect as seen in Fig.5.b, which are the units with a Δθ < 30° while the remaining units are sigmodal with a Δθ > 30°. We found units modulated only by reward or punishment in both NHPs and all cortical regions under study. At the same time, another segment of the population showed modulation by both simultaneously (Fig.7.a), which is a pattern seen in canonical affective brain regions such as the amygdala (Belova et al., 2007). Valance appears to be more strongly encoded in PMd, M1, and S1 compared with motivation. This is seen by the yellow regions in Fig.7.b for units that modulated significantly for both R and P. More units were significantly modulated during the post-reinforcement period for R only as seen in Fig.7.c compared to post-cue in Fig.7.b. In both NHPs, and in general, all brain regions showed more up modulated firing rates post-cue for units that were modulated by reward, and this trend was flipped for punishment modulated units, see Fig.8. Relations between post-cue and post-reinforcement for reward and punishment units were also flipped, respectively, with reward units going from up-modulated to down-modulated and punishment from down to up as one moves from post-cue to post-reinforcement. Finally, in Fig.11 and table 5, we see that most units in both NHPs and brain regions have more significant fits to model #4, which utilizes the actual R values from the task and only considers R values in the model, however not as clearly for NHP P that encountered a more balanced distribution of R and P trials due to its higher error rate. Model #4’s prominence may be due to reward not only increasing the motivational aspect of the motor system, but perhaps also that reward leads to approach behaviors and increases neural firing rates compared to punishment units’ modulation, see Fig.8. For a trial to end in a punishment timeout period the NHP must have made an error during the trial. Thus, it is possible that the punishment trials are trials where the NHPs’ attention was not as high as it needed to have been for success. Further work is needed to test this attentional hypothesis.

### Towards affect agnostic Brain-Machine Interfacing (BMI)

Understanding neural modulation due to affect is relevant to biomedical engineers creating brain-computer interfaces (BMIs) with activity from sensorimotor regions (Sanchez et al., 2011; Marsh et al., 2015; An et al., 2018; Zhao et al., 2018; Atique and Francis, 2021). For BCI control signals to be accurately decoded from neural activity, it is important to determine if these units utilize a divisive normalization scheme for affective stimuli or naturally changing moods and emotional states. Recently the impact of context on decoding neural information has been addressed in the BMI literature when dealing with kinematics under BMI control switching from two dimensional to three-dimensional space (Rasmussen et al., 2017), as well as between reaching movements with and without expected object contact (Downey et al., 2017). Directional tuning in the M1 is modulated by reward during manual tasks (Ramakrishnan et al., 2017) and BMI control (Zhao et al., 2018). The latter study noted that grip force-related tuning functions are likewise modulated by reward expectation in M1. Affective information in the sensorimotor system could have profound implications for BMI-controlled robotic limbs, computer cursors for communications, and somatosensory neuroprosthetics as part of a bi-directional BMI (Choi et al., 2016; McNiel et al., 2016b, 2016a; Zhao et al., 2018; Kumaravelu et al., 2020; Quick et al., 2020). Affective information also influences correlational structure at the single-unit level (Moore and Francis, 2020), local field potential (LFP) level (An et al., 2018), and between these two (An et al., 2019). As LFPs can also be utilized for BMIs (Ince et al., 2010; Flint et al., 2013), affect’s influence on such neural activity is vital to understand in order to produce affect agnostic BMIs towards the restoration of movement control.

## Acknowledgments

Research was supported by NIH 1R01NS092894-01, NSF IIS-1527558.

## SUPPLEMENTARY INFORMATION

The figure below shows the electrode arrays’ placement in the two NHP subjects in the current study.

**Fig.S.1:**
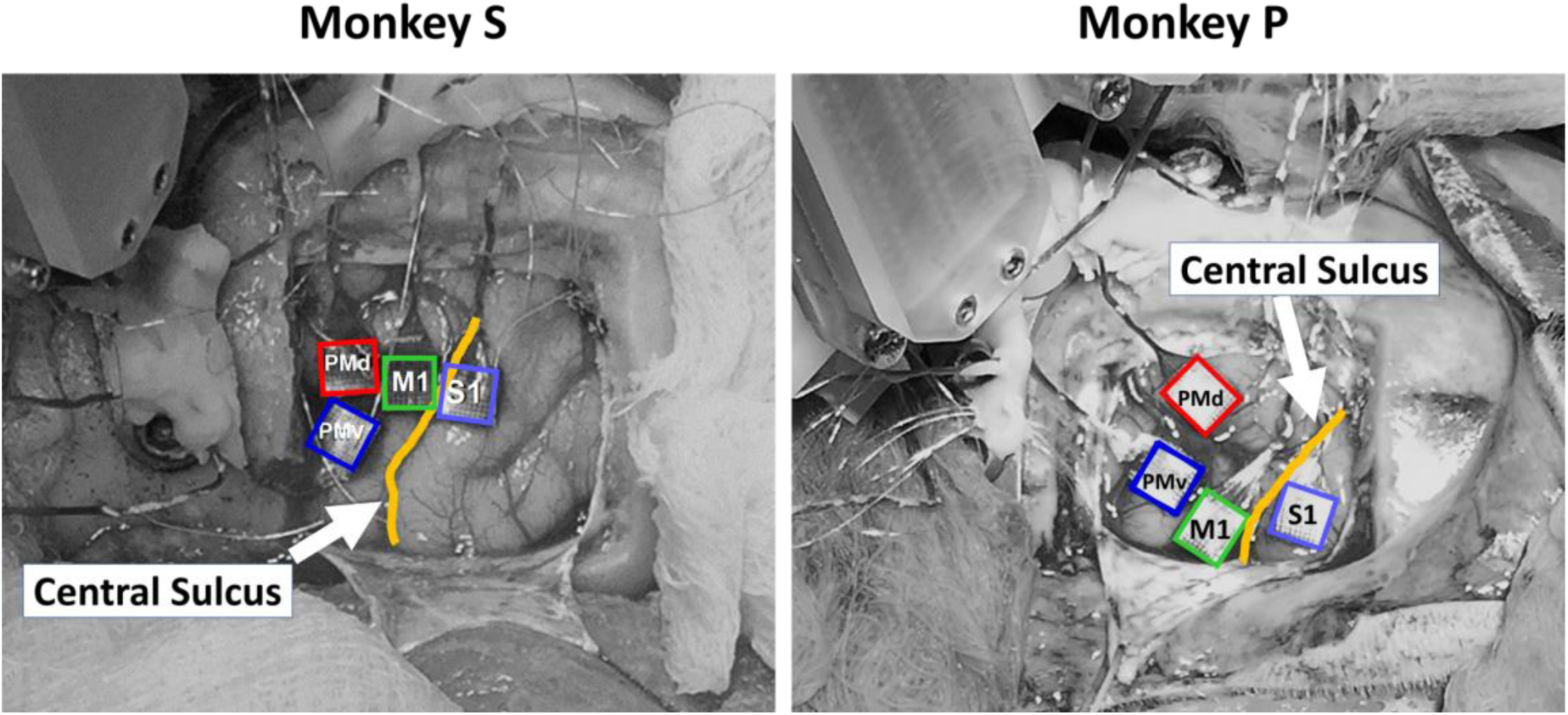
Position of four Utah arrays in relation to the central sulcus, yellow line, for NHP S (left) and P (right). The four arrays were implanted in caudal S1 (cS1), rostral M1 (rM1), PMd, and PMv cortices. Note that in NHP P, we had to implant the rM1 array more lateral than in NHP S due to a large set of blood vessels running through that region. Likewise, PMd was implanted more medial in NHP P for this exact reason compared to NHP S. PMv data was not utilized in this report due to noise issues.

**Fig.S.2.**
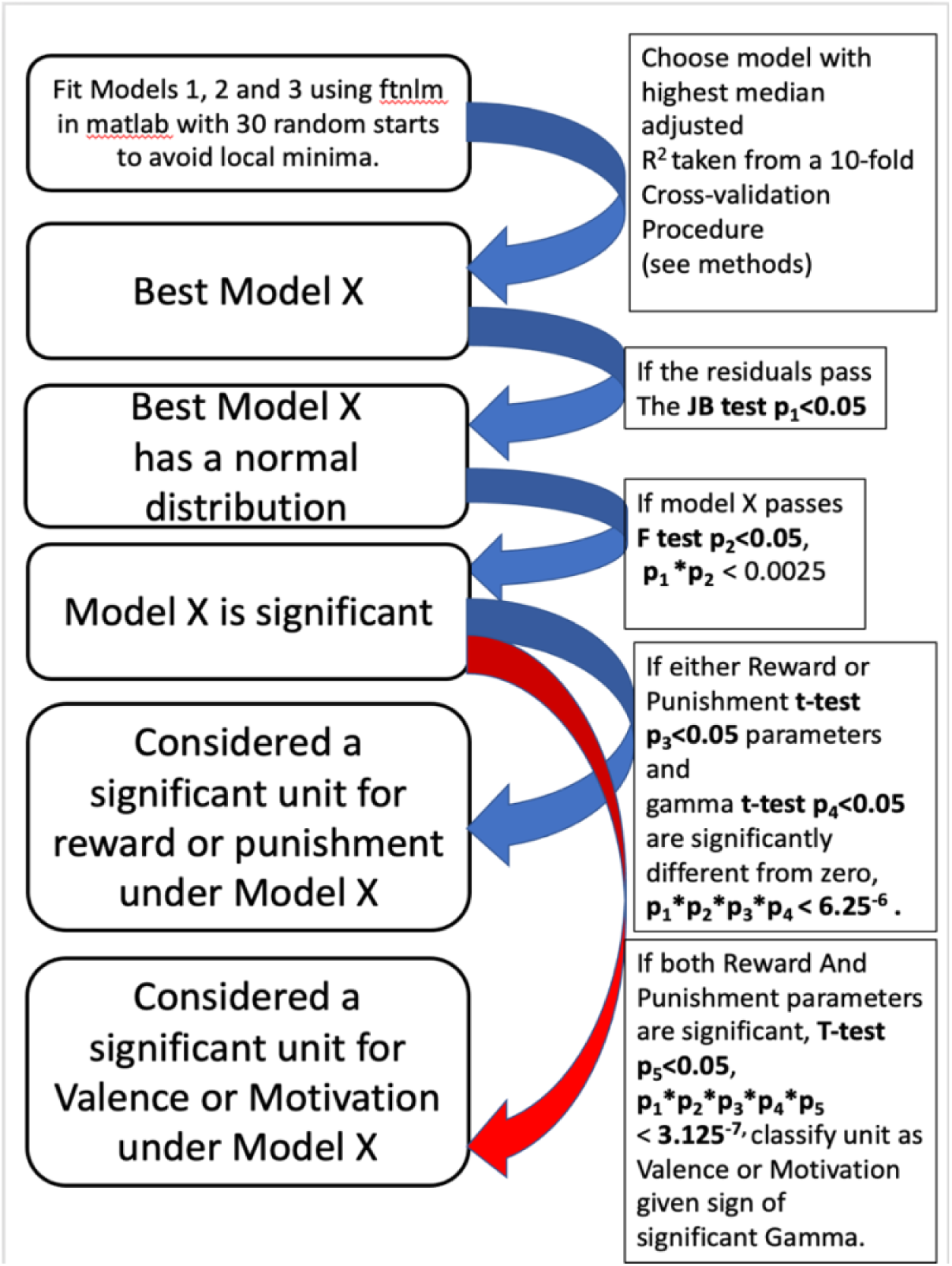
Successive Statistical Tests Give Increasing Statistical Significance. Our analysis divided the reward/ punishment modulation units into 3 groups: 1) Units only modulated by reward. 2) Units only modulated by punishment. 3) Units modulate by both reward and punishment. In group 3, we then analyzed if the units were valence or motivational (see figure.7b and 7c). As seen from Fig.S.2, we are not running these statistical tests in parallel but rather in succession. The associated p values given for the successive testing assume independence of tests and can be seen as the best-case p-value.

**Fig.S.3:**
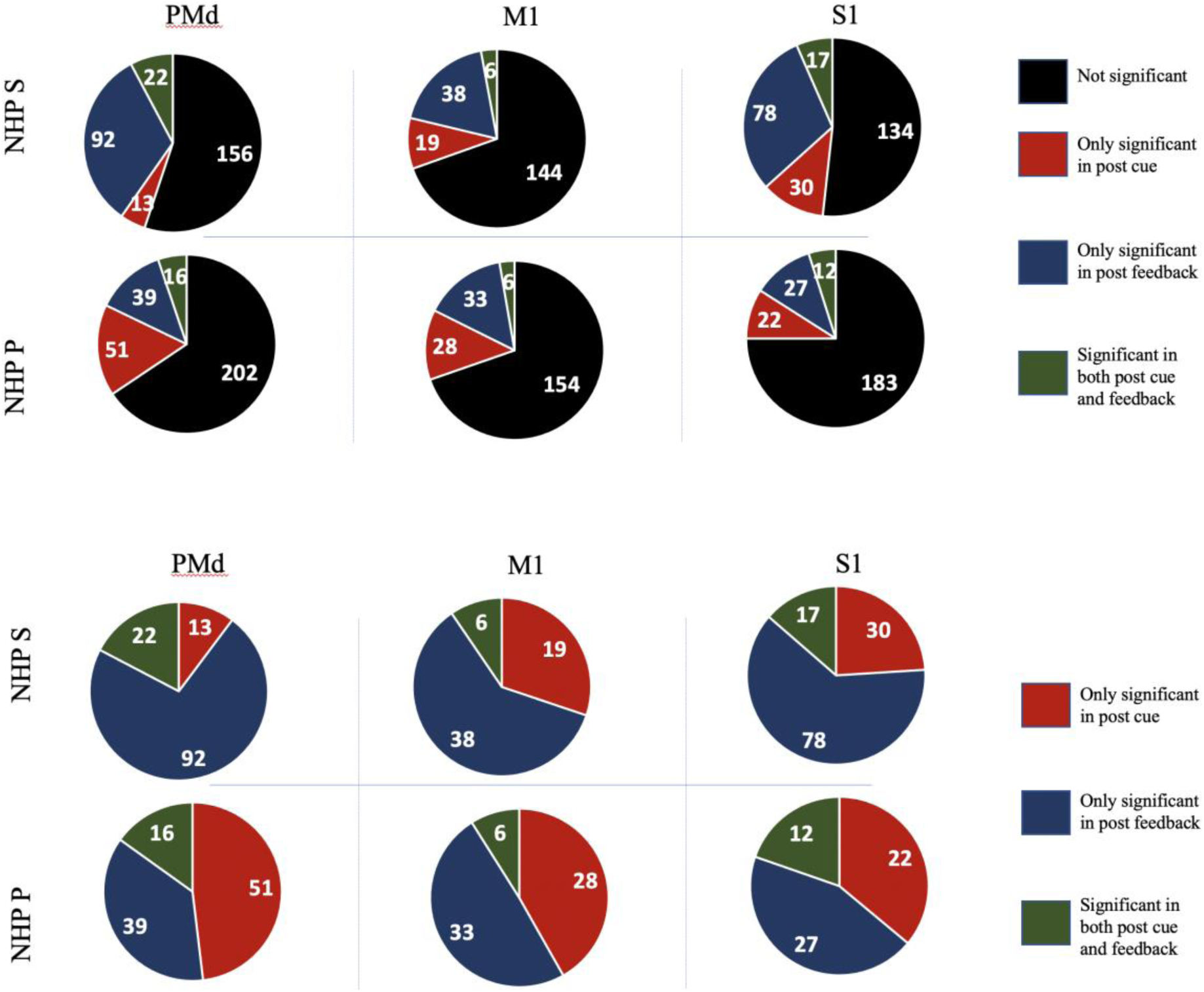
Here, we show the significant units and their makeup and the perspective of the larger non-significant population (Top subplot). For convenience, we have copied Fig.7 from the main text below it.

**Fig.S.4.**
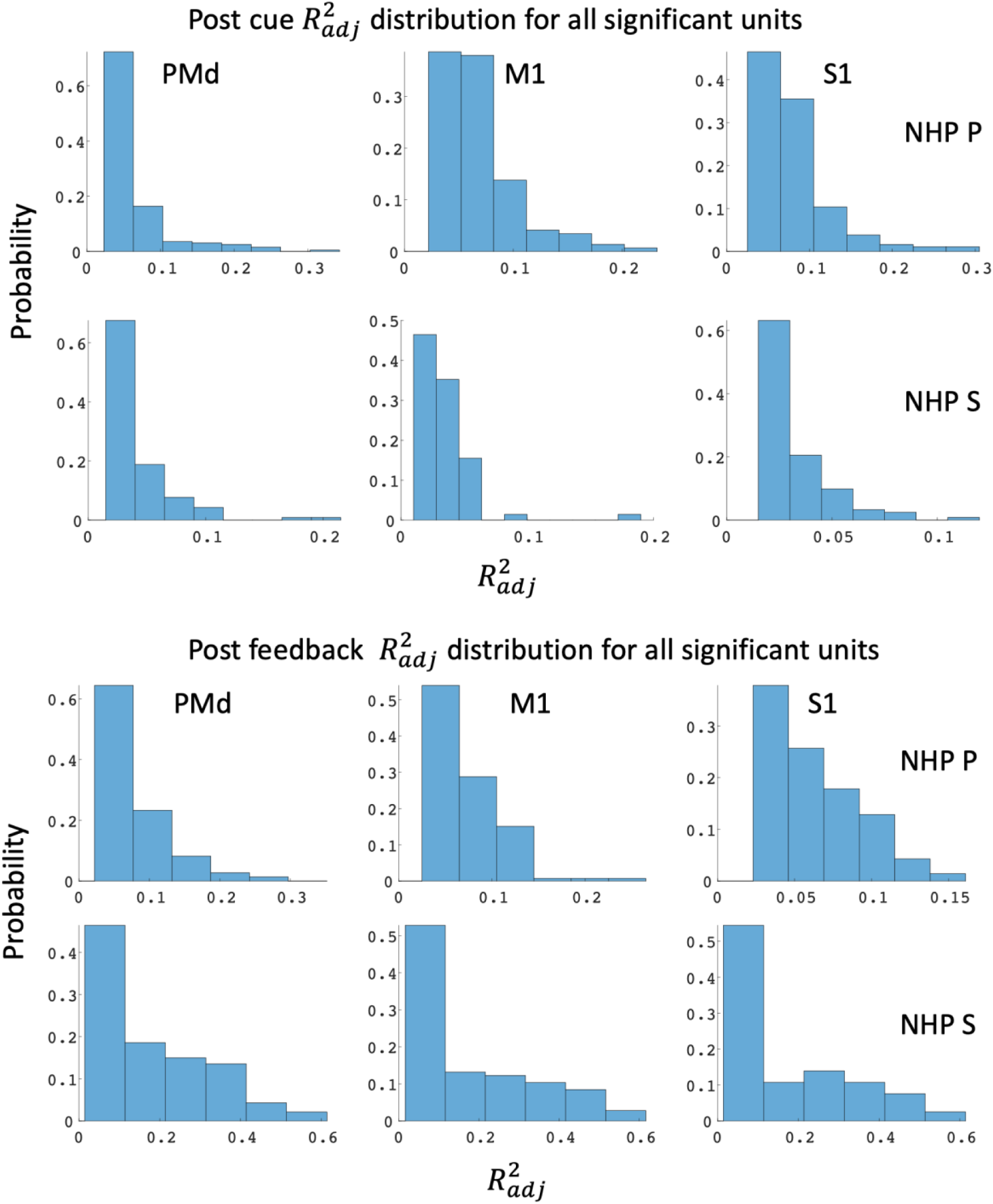
Distribution of the adjusted R squared values for the best significant model fit for a given unit during the post-cue, top two rows, and the post-feedback, bottom two rows, for the two NHPs labeled on the right-hand side of the figure. Fitting firing rates from every trial as compared to Fig.9 in the main text that comes from the model fits to the mean for each of the 16 different affective stimuli.

**Fig.S.5.**
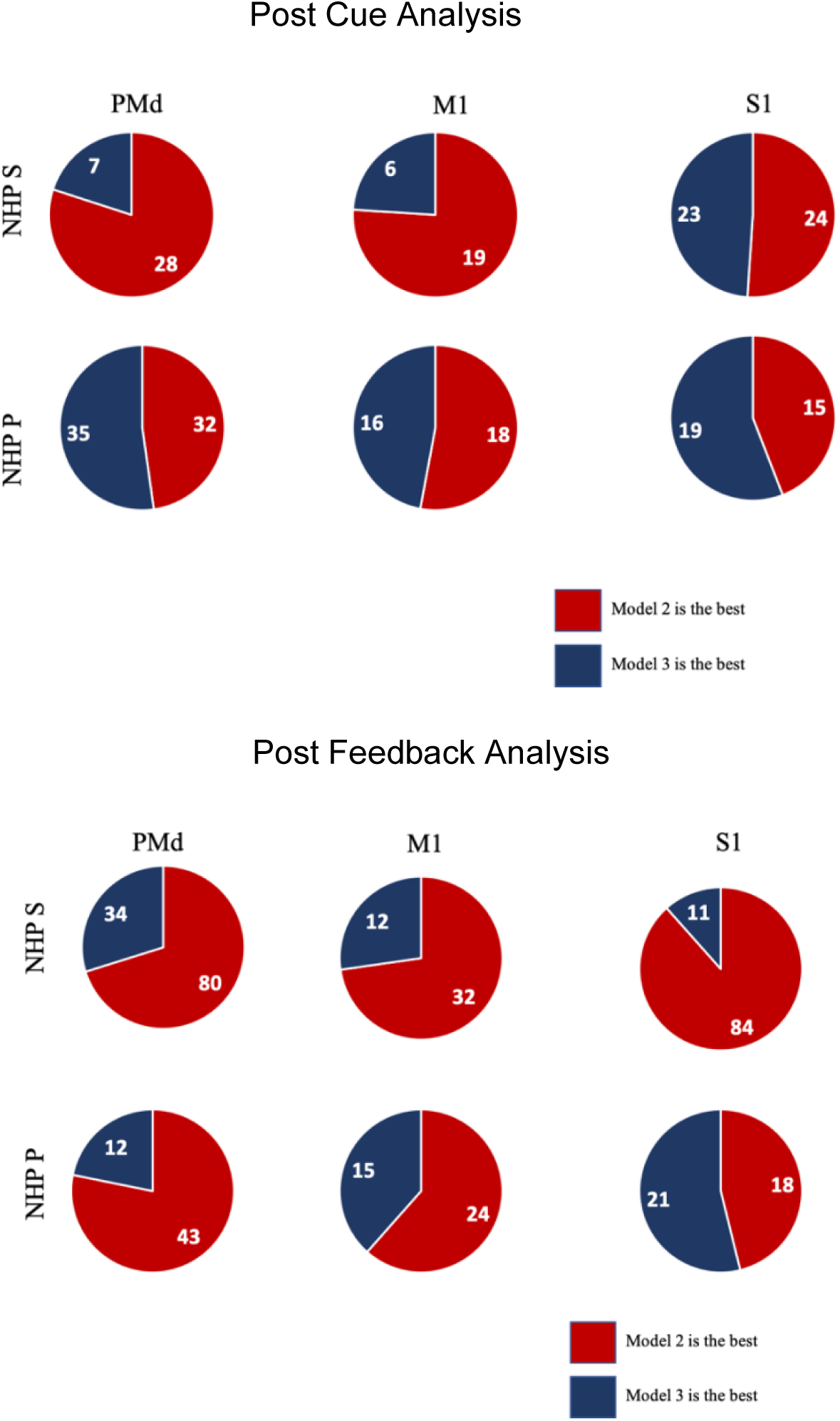
showing only significant units as compared to all units as seen in the main text table.3. Often, but not always, model #2, which utilized the R and P levels from the task in the denominator, outperformed model #3, which used the local population activity.

